# Intracellular Proteomics and Extracellular Vesiculomics as a Metric of Disease Recapitulation in 3D Bioprinted Aortic Valve Arrays

**DOI:** 10.1101/2023.06.22.546103

**Authors:** Cassandra L. Clift, Mark C. Blaser, Willem Gerrits, Mandy E. Turner, Abhijeet R. Sonawane, Tan Pham, Jason L. Andresen, Owen S. Fenton, Joshua M. Grolman, Fabrizio Buffolo, Frederick J. Schoen, Jesper Hjortnaes, Jochen D. Muehlschlegel, David J. Mooney, Masanori Aikawa, Sasha A. Singh, Robert Langer, Elena Aikawa

## Abstract

In calcific aortic valve disease (CAVD), mechanosensitive valvular cells respond to fibrosis- and calcification-induced tissue stiffening, further driving pathophysiology. No pharmacotherapeutics are available to treat CAVD, due to the lack of: 1) appropriate experimental models that recapitulate this complex environment; and 2) benchmarking novel engineered AV-model performance. We established a biomaterial-based CAVD model mimicking the biomechanics of the human AV disease-prone fibrosa layer, 3D-bioprinted into 96-well arrays. LC-MS/MS analyses probed the cellular proteome and vesiculome to compare the 3D-bioprinted model vs. traditional 2D monoculture, against human CAVD tissue. The 3D-bioprinted model highly recapitulated the CAVD cellular proteome (94% vs. 70% of 2D proteins). Integration of cellular/vesicular datasets identified known and novel proteins ubiquitous to AV calcification. This study explores how 2D vs. 3D-bioengineered systems recapitulate unique aspects of human disease, positions multi-omics as a novel technique for the evaluation of high throughput-based bioengineered model systems and potentiates future drug discovery.

## INTRODUCTION

Calcific aortic valve disease (CAVD) is an active, cellular-driven, progressive disease characterized by fibrotic valve thickening followed by leaflet calcification, valve stenosis, and ultimately heart failure and death^1–5^. Due in part to a lack of appropriate experimental models, no effective pharmacological intervention is available^6^. AV leaflets comprise three distinct layers defined by their extracellular matrix (ECM) composition^7,8^: the collagen-rich fibrosa layer, the proteoglycan-rich spongiosa layer, and the elastin-rich ventricularis. Valvular interstitial cells (VICs) are the most prevalent cell type in the AV. Under physiological conditions VICs are quiescent fibroblast-like cells and maintain valve homeostasis through proliferation and tissue remodeling^3,9–14^. However, under pathological conditions these VICs become activated^15,16^ and transform into myofibroblast-like cells or pro-calcific osteoblast-like cells that actively deposit hydroxyapatite in the ECM^17–19^. VICs embedded in the stiffer fibrosa layer drive calcification which spreads to the spongiosa, leaving the ventricularis relatively unaffected^20–22^. Therefore, the ability to study VICs within an environment like the disease-prone fibrosa is crucial to increase understanding of CAVD pathology.

Two-dimensional (2D) static VIC monoculture has been used extensively to study AV calcification^23–27^. In the context of CAVD, these models do not allow for the critical interaction between VICs and the ECM^6,28–33^, nor do they enable expansion towards co-culture models that incorporate interactions with inflammatory infiltrate and endothelial cells, known modulators of aortic stenosis^8^. To that end, three-dimensional (3D) hydrogel-based constructs have been used to create scaffolds that mimic the native AV architecture. Many different materials – synthetic polymeric and natural materials – have been used for 3D VIC culture. Each hydrogel material produces distinct biomechanical properties. Since VICs are mechanosensitive^53^, it is crucial that the properties of the hydrogel system are tailored to the specific tissue conditions present in normal human AVs^48,54^. Previous studies have shown that a mixture of 5% GelMA, 1% HAMA and 0.3% LAP photo-initiator results in a photo-crosslinkable hybrid hydrogel whose mechanical properties can be tuned to match the layer-specific stiffness of the native CAVD^21,28,48,55^. This hydrogel model is able to maintain VICs in a quiescent state, unlike 2D culture, while still allowing pathological differentiation^21,28,31,48^. CAVD is understood to be a multi-factorial process, with tissue stiffening, inflammation, mineral deposition, genetics, and epigenetics all having been implicated^8^. Additionally, extracellular vesicles (EVs) secreted from valve cells form critical nidi for calcification initiation in calcific valve pathology^15,56^. This factor has yet to be assessed when comparing similarities and recapitulation of *in vitro* models to CAVD pathology.

Here, we leverage cellular and extracellular vesicle proteomics to characterize the recapitulation of in vitro models of this disease comprehensively and holistically by assessing cellular- and EV-derived proteome-level alterations that take place as a function of static biomechanic properties. In addition, we develop, validate, and benchmark arrays of a novel 3D-bioengineered model system of CAVD pathogenesis that are compatible with high-throughput drug screening platforms.

## RESULTS

### High throughput bioprinting platform creates arrays of a biomechanically relevant CAVD model

Methacrylated gelatin and hyaluronic acid (GelMA/HAMA) were used to engineer a hydrogel system laden with primary human VICs, tuned to AV layer-specific (and disease-driving) biomechanics (**Fig. 1a, b**). Bioprinting parameters were optimized for compatibility with 96-well plate arrays suitable for high-throughput screening (**Fig. 1b, Supp. Methods**). Final hydrogel morphology was highly consistent between wells (**Fig. 1c**).

**Fig 1.**
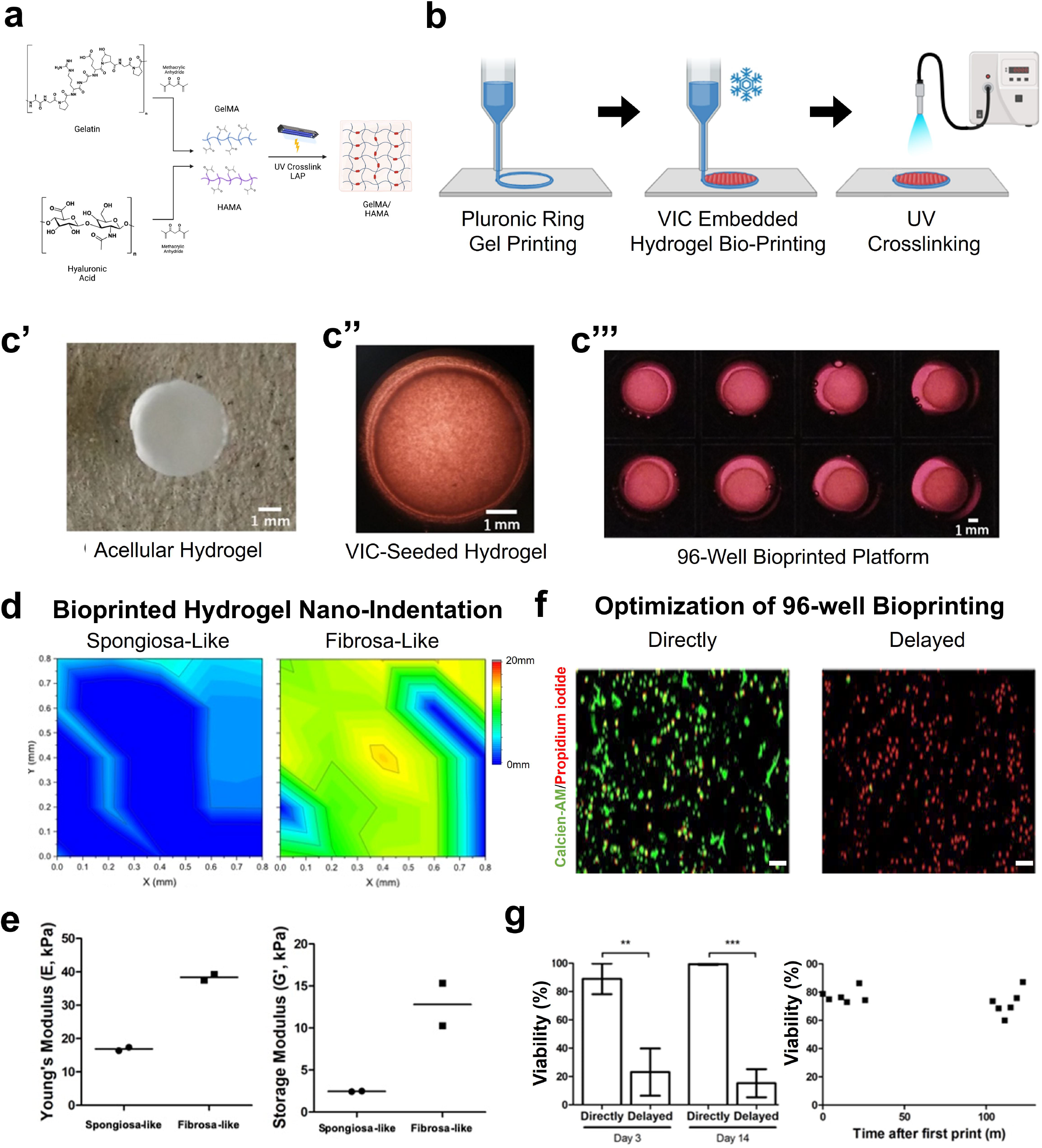

VICs sense and respond to stiffness of the ECM, therefore it is paramount for a successful CAVD model that material stiffness recapitulates that of the native AV layers ^45,48^. By modulating ultraviolet (UV) curing duration, the mechanical properties of the hydrogel were manipulated, and the mechanics of the spongiosa (disease-protected) and fibrosa (disease-prone) valve layer were replicated in a 96 well array format, based on previous studies and confirmed via nanoindentation analysis^21^ (**Fig. 1d,e**). Heatmaps of the nanoindentation measurements across the hydrogel surface showed spatially resolved consistency in biomechanical measurements (**Fig. 1d**) and were quantified to recapitulate known layer-specific biomechanics (**Fig. 1e**). Since calcification predominantly occurs in the fibrosa layer of the AV^20,21^, the fibrosa-like hydrogel model was used for subsequent experiments.

Cell viability was measured and optimized (**Fig. 1f,g**) to consider bioprinting-induced temperature, mechanical, and chemical stresses. Introduction of culture media into each well immediately after printing rescued otherwise-substantial decreases in cell viability (<25%) at both day 3 and day 14 post-printing that developed if medium administration was delayed until completion of the entire array printing. (**Fig. 1g, left**). Similarly, there was no significant difference in VIC viability between initial and final printed hydrogels (elapsed time = ∼2 hours), suggesting that cooling conditions and air-tight printing cartridge were able to slow cellular metabolism and reduce dehydration, maintaining viability (**Fig. 1g, right**).

Previous research has shown that calcification in VIC-encapsulated hydrogel models is significantly different between culture conditions after 14 days in culture^14,21,57^, thus informing our culture timeline. After 14 days in either normal media (NM) or two mediums with calcific stimuli: organic-phosphate osteogenic media (OM) or inorganic-phosphate pro-calcifying (PM), VIC viability within the fibrosa-like hydrogels stayed high across all media types and replicates (**Fig. 2a top, Fig. 2b left**). Valvular calcification can be the product of different processes like active mineral deposition by osteoblast-like VICs or that of apoptosis-related calcification, which is generally believed to be an artifactual process of *in vitro* culture^26,58^. We validated apoptosis was not inducing calcification via TUNEL staining, with showed little apoptosis-related cell death across all hydrogels and no significant differences between culture conditions, confirming the calcification is likely not mediated by cell death-related calcium accumulation (**Fig. 2a bottom, Fig. 2b right**).

**Fig 2.**
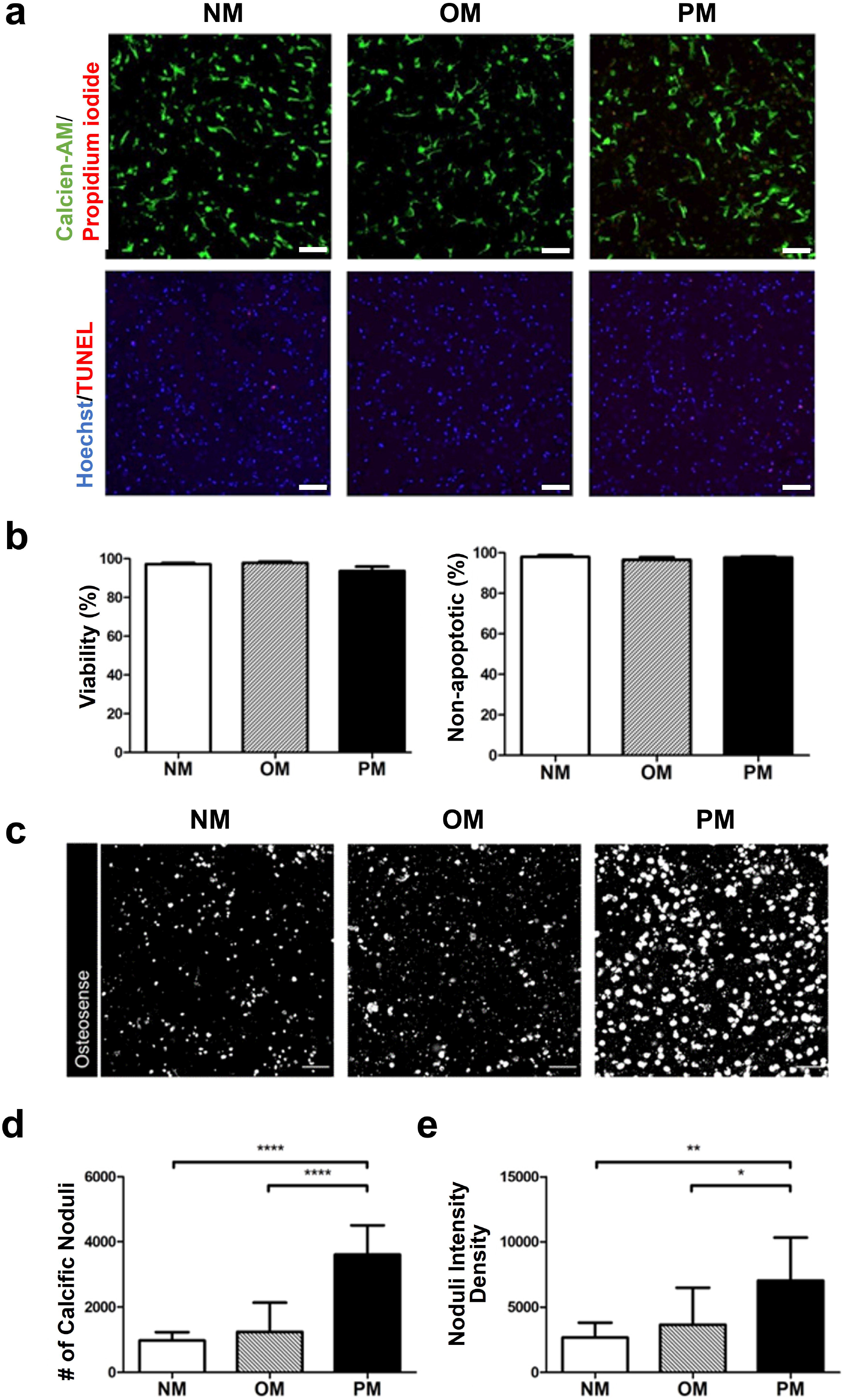

Calcification in the fibrosa-like 96-well hydrogel arrays was assessed by staining with the near-infrared calcium tracer (Osteosense680), which binds to hydroxyapatite crystals and detects microcalcifications^59^. All 3D-array media conditions showed some positive staining with Osteosense680 (**Fig. 2c**). However, trending increases and significant increases in both number of microcalcifications (**Fig. 2d**) as well as signal intensity (**Fig. 2e**) were seen in OM and PM conditions, respectively.

### Cellular proteomics identifies unique pathology modeled in 3D bioprinted arrays

Once viability and calcification induction were validated, proteomics techniques were used to assess how well these 3D bioprinted VIC hydrogel CAVD arrays (3D) recapitulated native CAVD tissue phenotypes (CAVD), compared to traditional 2D VIC monoculture conditions (**Fig. 3a**). Over 2,500 proteins were identified in 3D and 2D conditions, with >99% overlap of proteins identified between the two *in vitro* models (**Fig. 3b**). To remove potential background contaminants resulting from culture preparation, the proteome of the acellular hydrogel alone was also investigated and searched against *H. histolytica* (collagenase source), porcine (GelMA/HAMA source), bovine (culture-serum source), and human (to identify contaminant homology against target proteome) proteomes (**Fig. 3b**).

**Fig 3.**
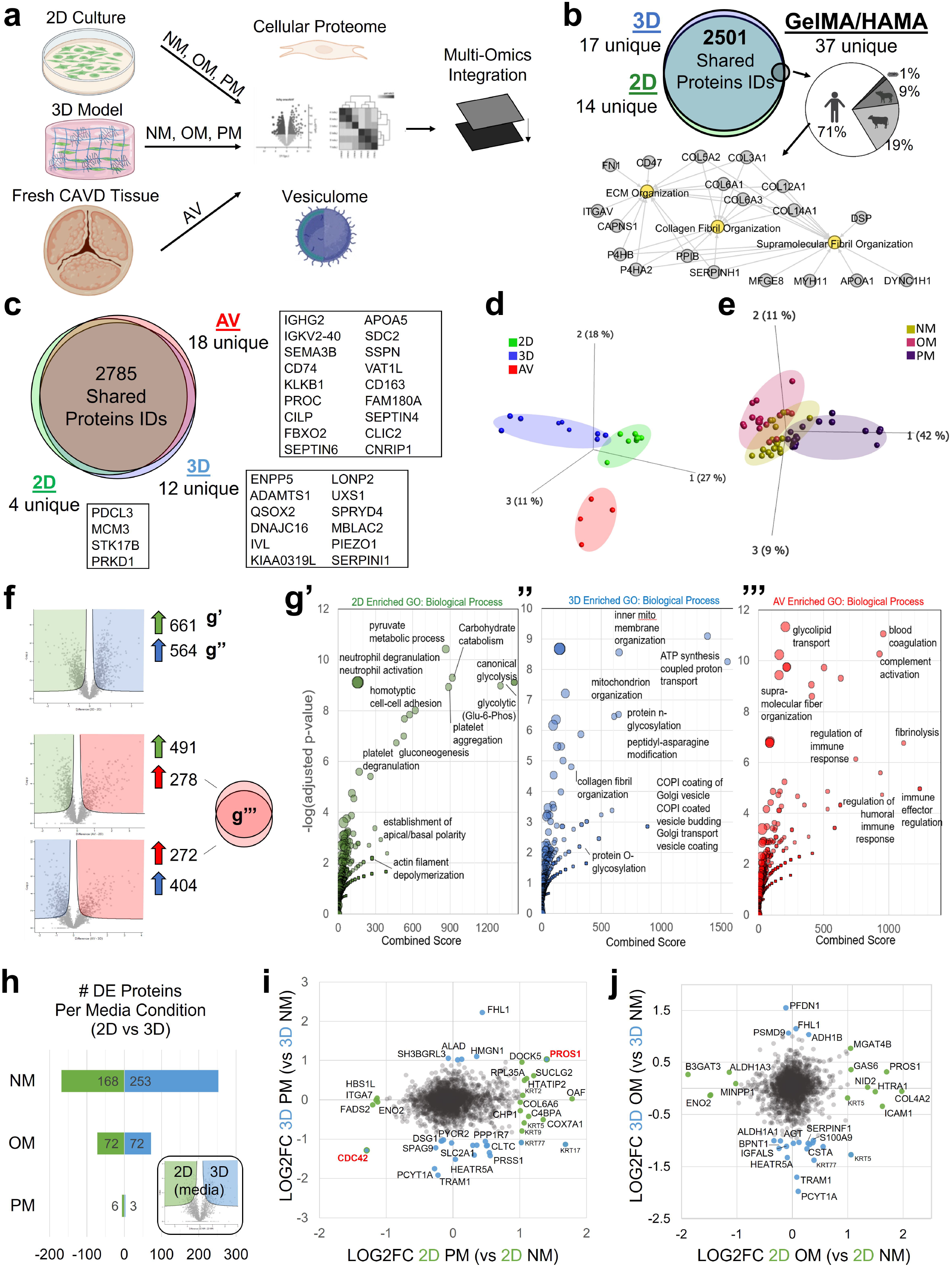

CAVD tissue-derived cells were included in subsequent analyses (**Fig. 3c**). Unfiltered principal component analysis (PCA) shows unbiased clustering of the three proteomes by model type (**Fig. 2d**) and by calcific media treatment (**Fig. 3e**, *in vitro* only). Between 2D and 3D *in vitro* models 48% of the measured proteome was differentially enriched (DE), while less than <30% of the proteome was significantly DE between CAVD and 2D or 3D – suggesting *in vitro* models’ cellular proteomes differ more from one another than they do from fresh tissue cellular proteomes (**Fig. 3f**). Using the DE proteins in **Fig. 3f**, we identified GO terms characterizing key differences between 2D and 3D conditions (**Fig. 3g’**,**g’’**) as well as both *in vitro* compared to tissue (**Fig. 3g’’’**). 2D cultures are significantly enriched in proteins related to platelet aggregation, homotypic cell-cell adhesion, canonical glycolysis, and actin filament depolymerization; while 3D cultures were enriched in proteins related to collagen fibril organization, protein N- and O-glycosylation, COPI coated vesicle trafficking, and mitochondrial organization. Finally, isolated CAVD cells were enriched in proteins related to regulating immune response, complement activation, and glycolipid transport (**Fig. 3g**) – consistent with the immune cell infiltration present in native CAVD.

To test the hypothesis that culture models responded differently to calcification induction media, the two *in vitro* datasets were analyzed by media condition and calcific stimuli, independent of the AV tissue dataset (**Fig. 3h-j, Supp. Fig. 1**). The NM condition showed the greatest amount of DE proteins (15%; 421/2,815), with OM having only 5% DE proteins (144), and PM only <1% (**Fig. 3h**). To compare how each *in vitro* model uniquely responds to calcific media, log_2_FC scatterplots of significantly DE proteins (-log_10_p>1) were plotted for both PM and OM. We found that PM (**Fig. 3i**) and OM (**Fig 3j**) media elicited distinct cellular responses in both *in vitro* models. In 3D PM media, enriched proteins were related to ECM assembly (AGT) and hyaluronan biosynthesis (CLTC), while 2D PM showed enrichment in proteins associated with smooth muscle cell contraction (DOCK5) and phospholipid biosynthesis (CHP1) (**Fig. 3i**). In OM media, 2D culture showed enrichment in two vitamin K dependent proteins (GAS6, PROS1) and a depletion in ENO2; while, 3D culture showed DE in cell growth regulators (S100A9, FHL1), and ROS metabolism (ADH1B, ALH1A1) (**Fig. 3j**). Interestingly, the only two proteins consistently DE in PM media were CDC42 and PROS1 – showing that each *in vitro* model has distinct responses to calcification induction. (**Fig. 3i**).

We then aimed to correlate *in vitro-*to CAVD-derived cellular protein abundances. Pairwise correlation analysis shows all *in vitro* media conditions were highly correlated to one another (r>0.9). While proteome-wide correlation analysis showed significant association of both 2D and 3D cells to CAVD cells (2D r_avg_=0.82; 3D r_avg_=0.79), individual protein abundance analysis must also be considered^60^. 2D and 3D protein abundances were compared against the AV dataset across all media conditions (**Fig. 4b,c; Supp. Fig. 2**). Within the 2D model, NM media had the greatest number DE proteins compared to native CAVD, with PM media being the most similar condition **(Fig. 4b)**; dissimilarly, 3D arrays showed few cell-derived proteins with DE to CAVD (**Fig. 4c**). Overall, the 3D model recapitulated the CAVD cellular protein abundance profile across 94% of proteins measured, while the 2D model recapitulated 70% of protein abundances.

**Fig 4.**
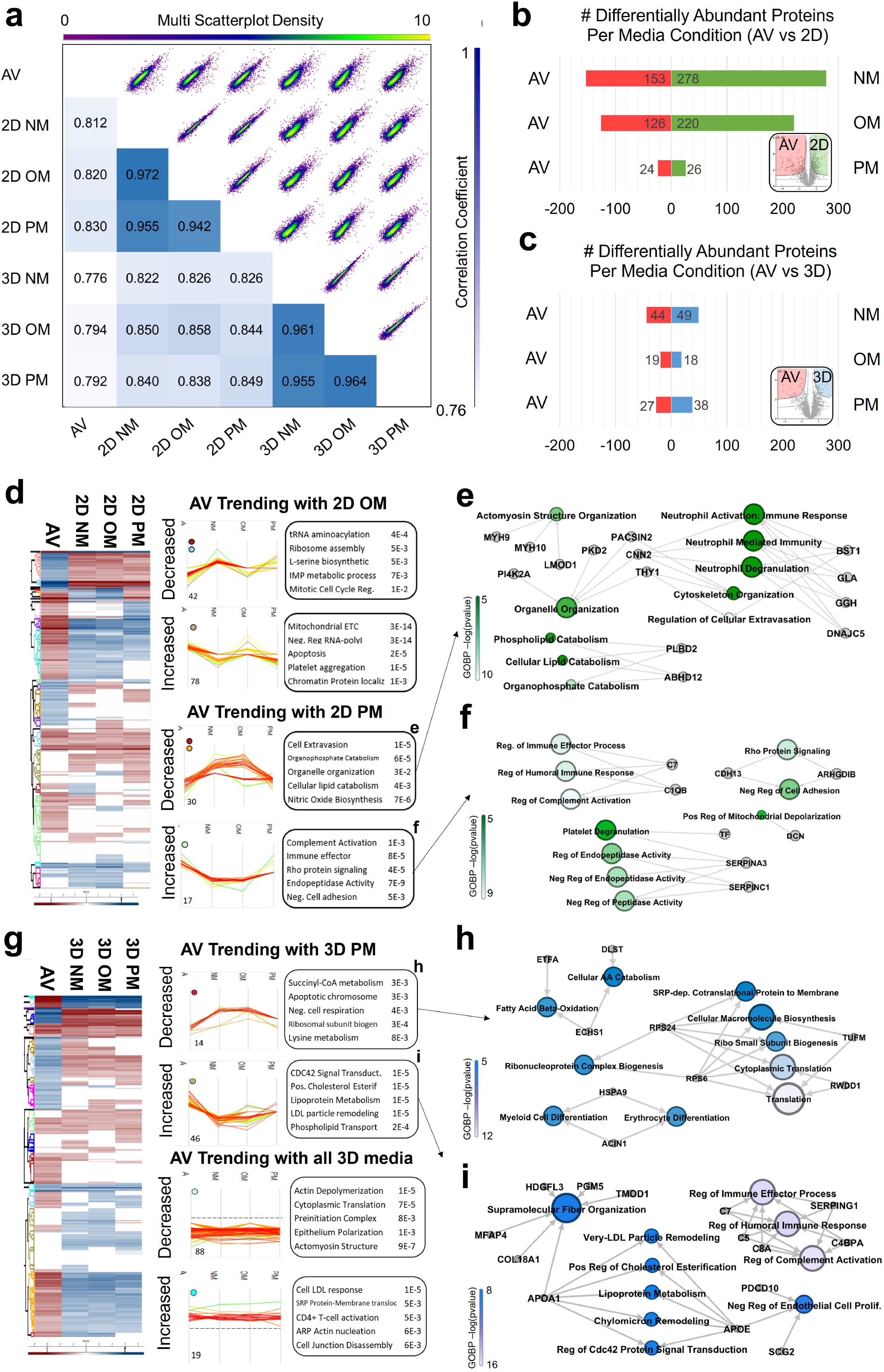

Next, we used protein trend analysis to identify which key proteins in 2D and 3D conditions best recapitulate the CAVD cellular proteome (**Fig. 4d-h, Supp. Table 2**). In the 2D PM condition, proteins recapitulating CAVD abundances were associated with cell adhesion (CDH13, ARHGDIB) and cytoskeletal organization (PACSIN2, CNN2, THY1) (**Fig. 4d-f**). In the 3D condition, both OM and PM medias recapitulated CAVD protein abundances associated with supramolecular fiber organization (MFAP4, COL18A1, TMOD1) lipoprotein metabolism (APOA1, APOE), negative regulation of endothelial cell proliferation (SCG2, PDCD10) and fatty acid beta-oxidation (ATFA, ECHS1) (**Fig. 4g-i**). We also identified a group of proteins that trended with CAVD tissue in all 3D media conditions (**Fig. 4g, bottom**). This cellular proteome centered analysis unbiasedly shows that the 3D hydrogel arrays recapitulate unique aspects of disease not captured in 2D monocultures.

### Extracellular vesicle cargo proteomics identifies differential loading in CAVD pathology

Literature shows a fundamental role for EVs in calcific cardiovascular disease, both in atherosclerosis and CAVD^15,56,61,62^. To evaluate the role of EVs in in vitro CAVD modeling, EV cargo proteomics was performed. EVs were isolated and an appropriate size range was quantified via nanoparticle tracking analysis (**Supp. Fig. 3**). Proteomics performed on isolated EVs identified over 1,300 proteins, including 26 common EV markers (**Fig. 5a**). Unbiased PCA analysis shows distinct clustering of model- and media-tagged EV proteomes (**Fig. 5b-c**). Unlike the cellular proteomics analysis, all media conditions (NM, OM, PM) resulted in a similar number of DE EV cargo proteins between *in vitro* models (**Fig. 5e**). However, the EV proteome was more stable overall: only an average of 5% (42/977) of the total proteome was DE amongst any media treatment between *in vitro* models (**Fig. 5e**), compared to 15% of the cellular proteome in NM conditions (**Fig. 3h**).

**Fig 5.**
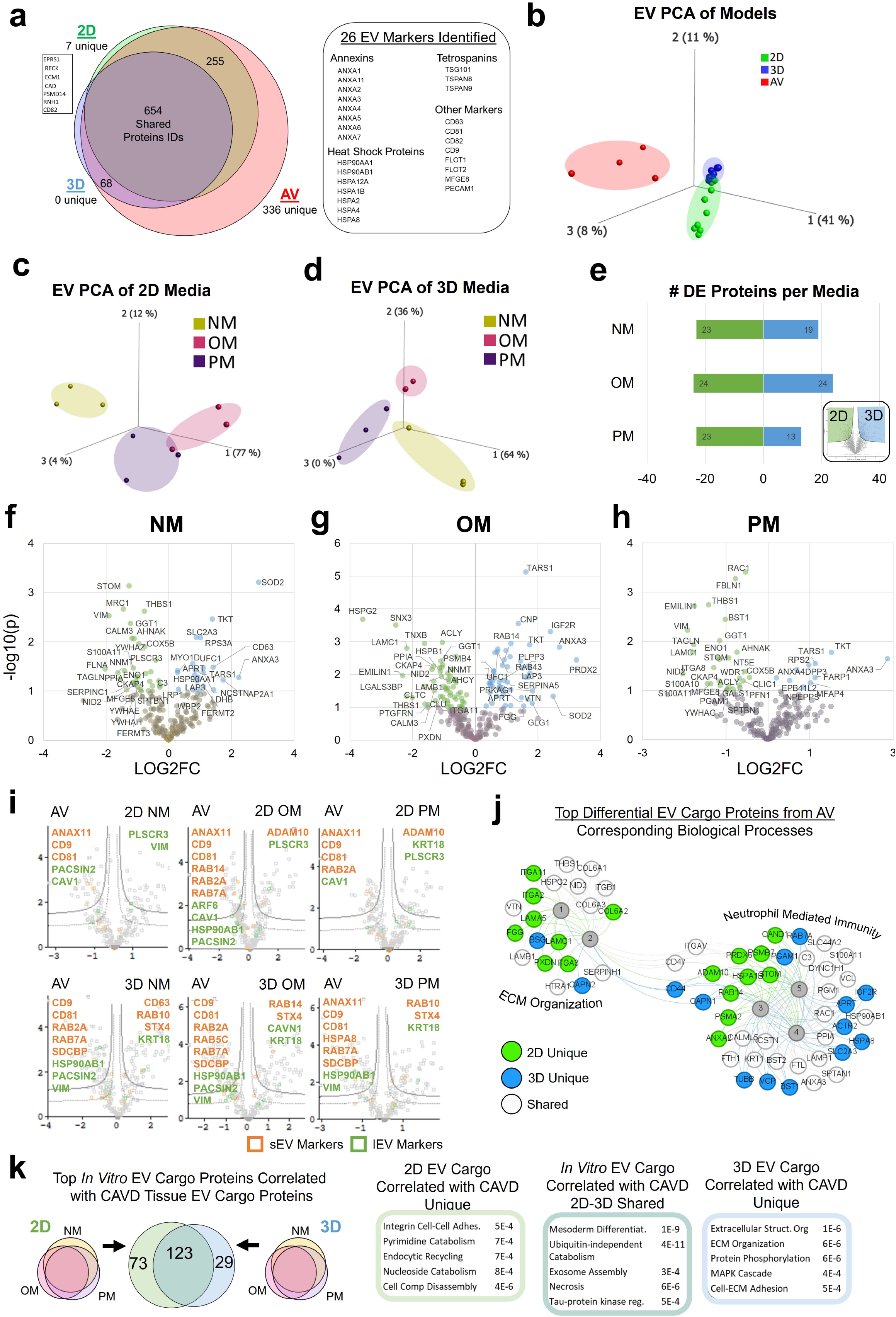

In the 2D vs. 3D models, less than 4% of all DE EV cargo proteins were shared between media treatments, suggesting a unique loading response to each media and model (**Fig. 5f-h**; **Supp. Fig. 4**). Key *in vitro* model EV cargo differences in NM media were related to integrin mediated signaling (CD63, FLNA, FERMT2, FERMT3) and mitochondrial membrane dependent apoptosis (YWHA-family) (**Fig. 5f**). OM EV cargo differences were related to superoxide regulation (PRDX2, APOA4, SOD2) and ECM assembly (TNXB, PXDN, LAMB1, EMILIN1) (**Fig. 5g**). Finally, alterations to PM EV cargoes were related to plasminogen^63^ (ENO1, THBS1) and, again, ECM (VIM, MFAP4, EMILIN1) (**Fig. 5h**).

We identified that AV EV cargo proteomics were enriched in many markers of both small and large EVs (**Fig. 5i**). From the differential enrichment analysis in **Fig. 5i**, we identified proteins that were not well recapitulated in CAVD tissue EV cargo in both *in vitro* models (**Fig. 5j**). The DE proteins from both *in vitro* models were best described by two overarching categories – ECM organization (informed primarily by 2D unique DE proteins), and neutrophil mediated immunity (informed by 2D and 3D DE proteins) (**Fig. 5j**).

Finally, we aimed to determine which *in vitro* model EV cargoes were best recapitulating disease. Here, we identified EV cargo proteins ubiquitously identified in AV and calcific models (2D, 3D, and CAVD) were related to exosome assembly (SDCBP, PDCD6IP), ubiquitin-independent catabolism (PSM family), and mesoderm differentiation (integrin family). 2D and CAVD shared EV proteins were related to integrin cell-cell adhesion (DPP4, PODXL, PLPP3) as well as cell composition disassembly (BSG, CAPN1-2, HTRA1, CD44). As expected, 3D and CAVD shared EV cargo proteins with correlated abundance profiles were related to ECM organization and structure (ITGA11, COL6A3, LAMC1, NID2, HSPG2) as well as cell-ECM adhesion and MAPK cascade and phosphorylation (CAV1, PBLN1, EMILIN1) (**Fig. 5k, Supp. Table 3**; **Supp. Fig 5**). Overall, these EV results are consistent with our findings in the cellular proteomes and demonstrate the importance of careful *in vitro* model design/selection when attempting to appropriately recapitulate specific aspects of cellular responses to disease-driving biomechanics, matrix material properties, and inter/intracellular signaling cascades with high-fidelity to those of human tissues.

### Integration of multi-layer omics datasets shows key drivers of CAVD recapitulation

After deeply characterizing differential abundance patterns in cellular and EV proteomes, we aimed to study patterns of associations between these proteomes to identify novel proteins that recapitulate the CAVD pathology within 3D printed hydrogel model and 2D culture condition under different stimuli. For this, we leveraged two computational analyses methods: LIONESS and rCCA, each elucidating different aspects of associations (**Supp. Fig 6**).

Integrative analysis of cellular and EV proteome was done using LIONESS^64–66^. In this approach, we computed principal components (PCs) that explain 95% of the variance of each dataset (Cell: cPC, EV: ePC) (**Supp. Fig 6a-b**) and used these as nodes in a correlation network bipartite graph between the reduced dimensions of proteome. The loadings of each PCs represent a pattern of highly correlated proteins in the two layers. To further understand these correlation patterns between different calcification models with NM control, we construct sample-specific networks^64,65^. These sample specific networks allow us to identify most significantly different PC pairs and their loadings, highlighting each protein’s ranked contribution to variation between calcific and normal media treatments.

For each of the four calcifying comparisons (2D OM, 2D PM, 3D OM, and 3D PM vs. respective NM), we chose the top 2 PCs along with their loadings and extracted the top 20 proteins in each and represented them into a composite network.(**Fig. 6a**). The network comprises three types of nodes: compared calcification model (cases, rounded squares), associated two PCs for cell and EV layer (diamonds), and proteins from loadings of those PCs (circles). The color of the nodes indicates the proteome layer they belong to, with yellow representing the Cellular layer and pink representing the EV layer. The protein nodes are further categorized based on their sharedness across models, and the network is organized from the most shared to the most specific (**Fig 6a, tiers 4-1**, respectively).

**Fig 6.**
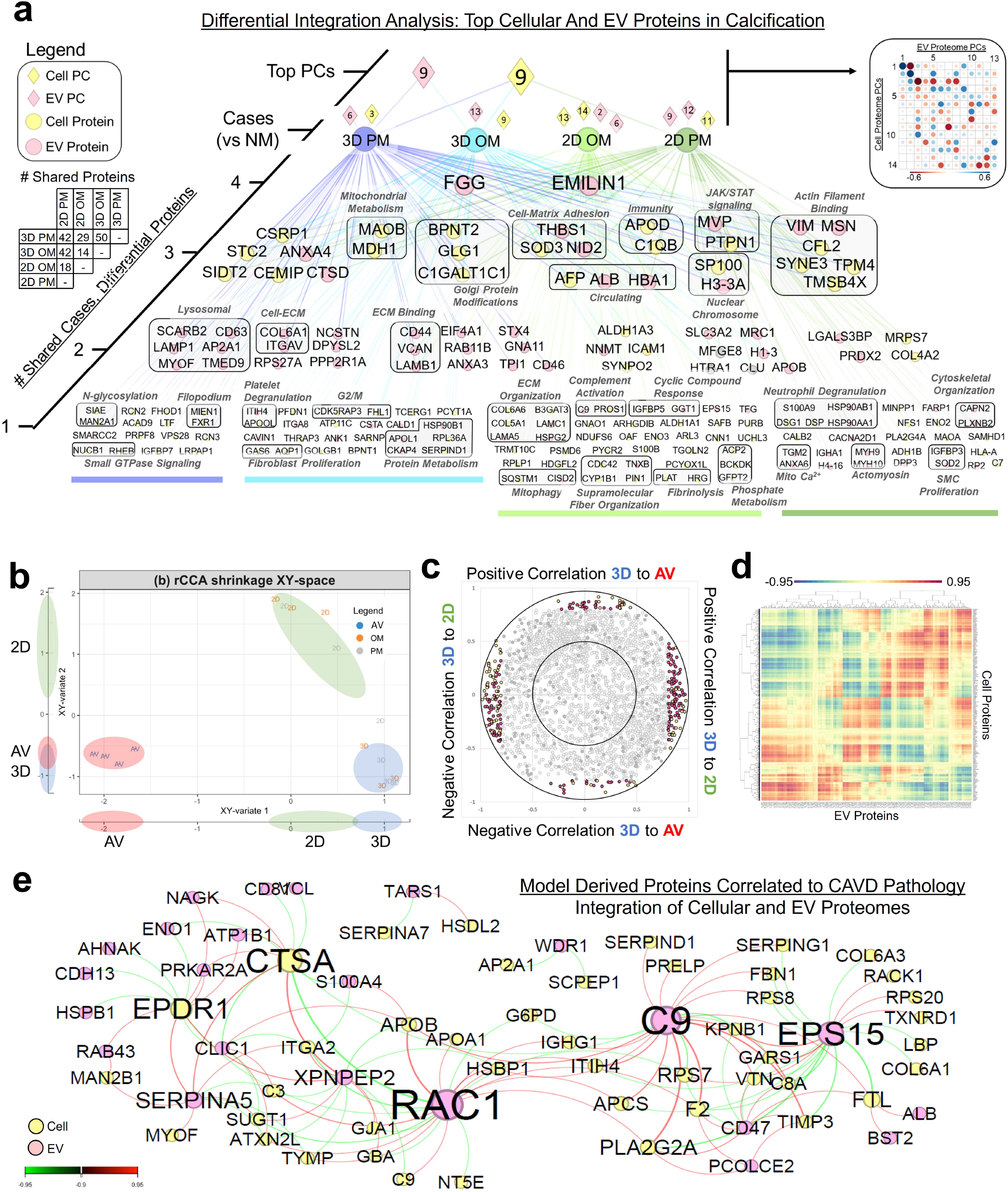

Over 60 cell- and EV-derived proteins were shared with at least one other condition (**Fig. 6a, tiers 2-4**), highlighting proteins that drive calcification in CAVD models independent of dimensionality or phosphate type. Interestingly, only four proteins identified in this integrated analysis to drive calcification were shared between the cellular and EV proteomes – vitronectin (VTN), lactadherin (MFGE8), clustering (CLU), and calprotectin L1H (S100A9). Only two proteins were associated with all four conditions: fibrinogen gamma (FGG) and elastin microfibril interfacer (EMILIN1), both as EV-derived proteins. Within unbiased subclusters, proteins of shared biological processes are grouped (**Fig. 6a**). Amongst cellular and EV proteins shared between 3 conditions, processes highlighted include mitochondrial metabolism (MAOB, MDH1), cell-matrix adhesion (THBS1, SOD3, NID2) and actin filament binding (VIM, MSN, CFL2, SYNE3, TPM4, TMSB4X). Interestingly, only 19% (41/215, p < 0.05 from exact binomial test) of proteins identified have been previously implicated in valve disease (PheGenI, NCBI). This integrated analysis highlights known and novel drivers of calcification *in vitro*.

Finally, we aimed to examine which proteins within both cell and EV datasets were driving recapitulation of calcified *in vitro* models and native CAVD tissue. We utilized regularized canonical correlation analysis (rCCA) to identify the largest correlation between orthogonal components, or variates (**Fig. 6b-e**). rCCA is an unsupervised approach that focuses on maximizing the correlation between the two datasets (Cellular proteome and EV proteome, with only calcifying conditions included in the analysis). Canonical variate 1 shows high correlation between the two *in vitro* models, 2D and 3D, while the canonical variate 2 shows high correlation between 3D and CAVD tissue (**Fig. 6b**) The correlation circle plot is used to visualize relationship between canonical variates, where each point represents a protein from either the cellular or EV proteome (**Fig. 6c**). By thresholding this data, we identify proteins significantly correlated with each canonical variate. (**Fig. 6c**,**d**). Here, clusters of subsets of variables (proteins) of the same type of correlation are observed (**Fig. 6d**). Complementing these plots is the corresponding relevance network for significantly correlated cellular and EV cargo proteins driving correlation between calcifying hydrogels and CAVD tissue (**Fig. 6e**). We find that while EV proteins account for less than 10% of both datasets, they account for over 30% of proteins identified to drive calcification in CAVD and *in vitro* models. Importantly, 98% of proteins (64/65, p < 0.05 from exact binomial test) identified to be recapitulating calcifying models with CAVD tissue have previously been implicated in cardiovascular disease (PheGenI, NCBI). These integrative approaches to multi-layer proteomic analysis identified both known and novel drivers of calcification and identified how this 3D hydrogel model shows improved disease recapitulation of CAVD *in vitro*.

## DISCUSSION

We present the first -omics-driven assessment of a biomaterial model of CAVD. In addition, we describe a method for creating arrays of a biomechanically tuned hydrogel model of CAVD within a 96-well platform suitable for high-throughput target identification and drug screening platforms. Recent studies highlight that proteomics is a powerful tool for both screening for potential therapeutic targets as well as identifying mechanisms of action (MOA) during drug screening^6,67–70^. These reports include a proteomics-driven MOA study linked with a high-throughput 96-well platform, though only in short-term (24 hours) 2D cell line monoculture^67^. In contrast, our approach enables long-term (14-day) culture of primary human cells in a 3D microenvironment that recapitulates key disease drivers found in human tissues, and which is compatible with MOA drug screening.

In this study, we noted mild spontaneous calcification in NM conditions. This may be due to exposure to shear stress and pressure up to 75 bar during the bioprinting protocol, both of which have been shown to induce myofibroblast activation^71,72^. Additionally, the VICs used in this experiment were isolated from diseased human valves, and it is possible that this isolation contained a population of osteogenic myofibroblast-like cells as we previously demonstrated^73^. Despite this, we identified significant proteomic changes in both the cells and EV cargo within calcifying media treated conditions in the 3D culture when compared to NM controls, showing feasibility of this model.

While we found that our 3D hydrogel model recapitulated the ECM proteome, cell-ECM interactions, and mitochondrial metabolism regulating protein profiles of human CAVD tissues, both 2D and 3D models were lacking in immune/inflammatory responses identified in tissue. While immune co-culture studies are limited in the context of valve calcification, recent single-cell studies demonstrate the importance of cell-cell communication across cell types in calcific valve disease^13,19,74,75^. The model presented in this study provides a foundation which can be augmented to incorporate additional layers of complexity such as co-culture of other cell types, (patho)physiological cyclic shear/stretch, and additional layer-specific biomechanics. While our study focused on properties of the fibrosa and spongiosa, the elastin-rich and disease-protected ventricularis has been shown to drive unique protein signatures under culture conditions and likely contributes predominantly to tensile, not compressive, stress-based valvular physiological responses *in vivo*^7^.

This study utilized species-specificity of peptides identified via bottom-up proteomics to curate a background proteome of the ECM and other potential contaminants. In future studies, we aim to use proteomics techniques targeted to the ECM to identify mechanical responses to collagen sub-types, post-translational modifications, and cell-matrix interactions^76–79^. These techniques can also be used to assess cellular and EV responses to diverse matrix microenvironments used within hydrogels. This EV analysis could also be expanded to identify drivers of exosome vs. microvesicle secretion and corresponding proteomes^80^.

While GelMA/HAMA has the advantage of tunability for biomechanics, a focus of our current study/model, the cell-matrix interactions of gelatin differ from those to native collagen subtypes. Similarly, the role of non-fibrillar collagen subtypes has recently been implicated in fibrotic valve disease^77^, which may drive future biomaterial development. In summary, the current study highlights the critical importance in recapitulating the biomaterial-based microenvironment within *in vitro* models of valve disease, as we show a robust effect on cellular and extracellular vesicle proteomes that impact putative disease-driving biological processes.

## SUPPLEMENTAL METHODS

### VIC Isolation and cell culture

Calcified aortic valve leaflets were obtained from patients undergoing aortic valve replacement at the Brigham and Women’s Hospital (Boston, MA, USA) due to aortic valve calcification and/or stenosis. Aortic valve leaflet acquisition and utilization was done according to protocols approved by the Institutional Review Board (IRB protocol #2011P001703/PHS). In this study, valves from ten individual donors were used. After resection, the valves were kept in Dulbecco’s Modified Eagle’s Medium (DMEM) for a maximum of 1 hour at 4 °C prior to digestion. Every leaflet was rinsed briefly in phosphate buffered saline (PBS), the surface of the valve was gently scraped with a razor blade and rinsed again. Next, the leaflet was cut up into small pieces with a razor blade and digested for 1 hour at 37 °C in 1 mg/ml collagenase (Sigma-Aldrich, St. Louis, MO, USA) that had been sterile filtered through a 0.2 μm syringe filter (Pall Life Sciences, Ann Arbor, MI, USA). The solution was mixed every 20 minutes and vortexed briefly after 1 hour. Afterwards, the supernatant was aspirated, the remaining tissue was washed in fresh DMEM, and fresh sterile filtered collagen was added to the tissue. The tissue was then incubated at 37 °C for 3 hours and was mixed every 30 minutes and vortexed briefly after 3 hours. Once finished, the digested leaflet suspension was passed through a 40 μm cell strainer (Thermo Fisher Scientific, Waltham, MA, USA) and the resulting solution was spun down at 1500 RPM for 5 minutes. Afterwards, the supernatant was aspirated, the pellet was resuspended in growth medium (10% fetal bovine serum (FBS), 1% Penicillin and Streptomycin (P/S)) and transferred to a Petri dish. The Petri dish seeded with the isolated VICs was incubated at 37 °C and 5% CO_2_. After two days, the cells were washed in PBS and fresh growth medium was added. The medium was changed twice a week until the VICs had passed 80% confluence. After this point, the medium was removed, and the cells were incubated in trypsin supplemented with ethylenediaminetetraacetic acid (EDTA) for 3 minutes at 37 °C. Afterwards, growth medium was added to neutralize the trypsin and the cells were spun down for 5 minutes at 1500 RPM. The supernatant was aspirated, the cells were resuspended in growth medium and split into new cultures. The cells were incubated at 37 °C and 5% CO_2_, with the growth medium replaced twice per week. VICs between passage 2 and 4 were used for bioprinting experiments.

### Methacrylated Gelatin (GelMA) production

Methacrylated gelatin (GelMA) was synthesized as previously described ^81^. Briefly, 20 g of gelatin from porcine skin (Sigma-Aldrich, St. Louis, MO, USA,) was suspended in 200 ml deionized water in a 500 ml flask which was stirred moderately for 1 hour. Next, the suspension was heated to 50 °C and stirred until the gelatin was completely dissolved. Once finished, 12.0 g methacrylic anhydride (Sigma-Aldrich) was added and stirred at 50 °C for 1.5 hours. Afterwards, the mixture was transferred to 50 ml conical tubes and centrifuged at 3500× g for 5 minutes. The supernatant was decanted and the remaining opaque solid left behind was diluted with two volumes of 40 °C deionized water. The mixture was transferred to dialysis tubing (10 kDa MWCO, SpectraPor 7, Spectrum Laboratories, Rancho Dominguez, CA, USA) and dialyzed against 3500 mL of deionized water at 40°C for 7 days. The water was changed twice a day. Afterwards, the content of the tubing was transferred to a beaker and using a 1 M NaHCO_3_ solution the pH was adjusted to 7.4. Next, using a 0.2 μm vacuum filtration unit and a polyethersulfone (PES) membrane, the solution was sterile filtered prior to transferring it into 50 mL conical tubes. Finally, it was snap frozen on liquid nitrogen and lyophilized until complete dryness. After approximately 10–14 days lyophilized GelMA as a solid white powder remained.

### Methacrylated Hyaluronic Acid (HAMA) production

Methacrylated hyaluronic acid was synthesized as previously described ^82^. Briefly, 1.0 g sodium hyaluronate (Lifecore Biomedical, Chaska, MN, USA) was dissolved in 100 ml PBS in a 250 ml flask and cooled to 4 °C. Next, 1 ml of methacrylic anhydride was added and stirred at 4 °C for 24 hours. A solution of 5 M NaOH was used to maintain a pH between 8.0 and 10.0. After 24 hours had passed, the solution was transferred to 50 ml conical tubes and centrifuged at 3500× g for 5 minutes. Afterwards, the supernatant was decanted into dialysis tubing and dialyzed against 3500 ml of deionized water at 4 °C for 7 days. The water was changed twice per day. Afterwards, the content of the tubing was transferred to a beaker and using a 1 M NaHCO_3_ solution the pH was adjusted to 7.4. Next, using a 0.2 μm vacuum filtration unit with a PES membrane, the solution was sterile filtered prior to transferring it into 50 mL conical tube. Finally, it was snap frozen on liquid nitrogen and lyophilized until complete dryness. After approximately 10–14 days lyophilized HAMA as a solid white powder remained.

### Bioink creation and VIC encapsulation

Stock solutions of 10 wt% GelMA, 3 wt% HAMA and 5 wt% LAP were made by dissolving lyophilized GelMA, lyophilized HAMA, and lithium phenyl-2,4,6- trimethylbenzoylphosphinate (LAP, Tokyo Chemical Industry Co., Portland, OR, USA) in 80 °C PBS. The pH of the HAMA solution was adjusted to 7.5 using 1 M HCl and the GelMA solution was sonicated for 1 hour at 37 °C. Afterwards, all solutions were heated to 80 °C and sterile filtered through a 0.2 μm syringe filter. The final stock solutions were stored at 4 °C until further use. Directly prior to printing these solutions were reheated to 37 °C. The 10 wt% GelMA, 3 wt% HAMA, and 5 wt% LAP stock solutions were mixed in DMEM at 37 °C to create the bioink with a 5% (v/v) GelMA, 1% (v/v) HAMA and 0.3% (v/v) LAP concentration.

Directly prior to bioprinting, the VICs were isolated from the culture flask by incubating them with trypsin supplemented with EDTA for 3 minutes at 37 °C. Afterwards, growth medium was added to neutralize the trypsin and the cells were spun down for 5 minutes at 1500 RPM. The supernatant was aspirated, and the pellet was resuspended in the bioink to create a bioink cell suspension with a final cell concentration of 10^7^ cells/ml of bioink. The bioink cell suspension was loaded into a clear plastic cartridge, the bottom and top were capped, the cartridge was wrapped in aluminum foil and placed on ice for 30 minutes.

### Toolhead extrusion pressure and temperature calibration

A temperature controlled toolhead (Cellink) containing a clear plastic cartridge with hydrogel was mounted to the first toolhead mount slot on the BioX 3D printer (Cellink, Cambridge, MA, USA) and a 22-gauge stainless steel needle (Cellink) were used for the calibration experiments. The temperature controlled toolhead was heated to 37 °C after which hydrogel extrusion was attempted, starting at 5 kPa and increasing stepwise with 5kPa increments until hydrogel extrusion was achieved. The minimal amount of pressure needed for extrusion was noted. After successful extrusion the temperature was lowered by 1 °C and after 5 minutes the minimal pressure needed for extrusion was recorded again until the setting of 7 °C was reached on the BioX. Additionally, it was noted at what temperature the hydrogel was no longer extruded as a liquid but as a more gel-like structure.

### Human VIC encapsulated hydrogel 3D bioprinting and culture

Designs were created in Tinkercad (AutoDesk, Inc., San Rafael, CA, USA) and exported as STL files which were imported into the BioX 3D Printer. The printer then spliced this file into a 3D code. The resulting Gcode file was then further edited with Sublime Text 3 (Sublime HQ, Pty Ltd., Darlinghurst, NSW, Australia). The BioX was used to print the constructs directly inside a black wall, clear bottom, 96-well plate (BRAND GMBH + CO. KG, Wertheim, Germany). A pneumatic toolhead (Cellink) containing a clear plastic cartridge with sacrificial pluronic gel (Allevi, Philadelphia, PA, USA) was mounted to the first toolhead mount slot and was used to print a cylindrical pluronic mold with a 22-gauge stainless steel needle (Cellink), at a pressure of 190 kPa with print speed of 6 mm/s and 100 ms preflow delay. The resulting mold had an outer diameter of 5.0 mm, inner diameter of 4.6 mm and total height of 1.5 mm. The temperature controlled toolhead (Cellink) was mounted to the second toolhead mount slot, set to 20 °C and loaded with the bioink cell suspension inside a clear plastic cartridge. The cartridge containing the bioink cell suspension was inserted into the 20 °C temperature controlled toolhead at least 30 minutes prior to printing. For a dual layer hydrogel, with a 22-gauge stainless steel needle, pressure of 75 kPa, print speed of 2 mm/s, a single layer of gelled bioink cell suspension was printed inside the pluronic mold. After extrusion the printer remained idle for 60 seconds to allow the bioink cell suspension to equally distribute inside the pluronic ring on the 37 °C printbed prior to UV crosslinking. The 365 nm UV toolhead (Cellink) was mounted to the third toolhead mount slot and UV light intensity was calibrated to 2.5 mW/cm^2^ using a radiometer (85009, Sper Scientific Direct, Scottsdale, AZ, USA). The first layer was crosslinked for 71 seconds. For all UV crosslinking the UV toolhead was positioned 1 mm above the top of the well. It was found that the highest UV intensity was recorded 2.3 mm behind the center of the UV toolhead. Therefore, this offset was manually implemented in all the UV crosslinking Gcodes. Next, from the same temperature controlled toolhead, the second bioink cell suspension layer was dispensed on top of the first layer and after another 60 seconds idling was crosslinked for 17 seconds by the UV toolhead. The bottom layer, the fibrosa like layer, was therefore crosslinked for 88 seconds in total, whereas the top layer, the spongiosa like layer, was crosslinked for 17 seconds.

For a single layer hydrogel model the preparation and setup is identical as the dual layer model. However now the Pluronic mold is filled up with the bioink cell suspension entirely, in a single layer. Afterwards, 60 seconds were allowed for the gel to melt again, and the entire hydrogel was crosslinked for 88 seconds. From this point onward both models were treated identically again.

Directly after printing of a hydrogel model had finished, 0.5 ml of growth medium was added to the well and all medium was replaced once more after all models were printed. One day after printing the growth medium was replaced with normal medium (NM; 5% FBS, 1% P/S), osteogenic medium (OM; NM, 10nM dexamethasone, 50μg/mL ascorbic acid, 10mM β-glycerolphosphate) or pro-calcifying medium (PM; NM, 50μg/mL ascorbic acid, 2mM sodium phosphate (NaH2PO4)) which was refreshed twice per week.

### Leaflet stiffness

To determine stiffness of the printed constructs, nanoindentation was performed on the hydrogel models. The Agilent G200 nanoindenter (Agilent, Santa Clara, CA, USA), equipped with a 90° diamond probe tip with a 50μm tip radius (DCMII, Micro Star Technologies, Huntsville, TX, USA) was used for mechanical testing. The tip area function was calibrated using a fused silica sample. Additionally, at a 5 μm pre-compression depth, a punch diameter of 45.153 μm was calibrated. The nanoindentation experiments were run as dynamic indentations with a frequency of 110 Hz to allow for the complex shear modulus of soft materials like the hydrogel models ^21,83^. Indentations were performed at room temperature while keeping the hydrogel constructs moist by means of an air humidifier. The energy stored during one oscillatory cycle was reported as the storage modulus G’, whereas the dispersed energy during one oscillatory cycle was reported as the loss modulus G”. The loss tangent was calculated as the ratio between G” and G’. The complex modulus G* was calculated by:

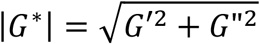

A Poisson’s ratio of 0.45 was assumed for the hydrogel construct based on previous indentation experiments ^21^. Young’s modules (E) was calculated for all printed models using:

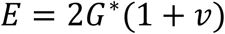

Nanoindentation was performed at 25 different spots, 200 μm apart on a 5 x 5 grid on each printed construct. Negative values were omitted, and the mean of the remaining measurements was calculated.

### Calcification, apoptosis, and live/dead assays

To determine the amount of calcification after 14 days in NM, OM or PM, samples were stained with the near infrared fluorescence agent OsteoSense 680EX (PerkinElmer, Waltham,MA, USA). Prior to imaging, all hydrogels were incubated overnight with the imaging agent at a 1:100 dilution in NM at 37 °C and 5% CO_2_. To assess VIC viability after printing, the cell laden hydrogels were stained with the double staining live/Dead kit (Sigma-Aldrich, St. Louis, MO, USA). The hydrogel models were first washed in PBS prior to 30 minutes of incubation at 37 °C, 5% CO_2_ with the fluorescent imaging agents which consisted of a mixture of 40μM Calcein AM and 20μM propidium iodide. Apoptosis was assessed with the Click-iT TUNEL Alexa Fluor 594 kit (Thermo Fisher Scientific, Waltham, MA, USA) according to the manufacturer’s instructions. A Nikon A1 confocal microscope (Nikon Instruments Inc., Melville, NY) with NIS Elements (version 5.10) and Zeiss LSM 880 with Zen (blue edition) were used to capture z-stacks at 10 μm per slice per gel construct. Image J was used to compress 10 slices of the z-stacks into maximum intensity projections, remove the background noise and measure signal intensity and number of positive signals.

### Hydrogel digestion, VIC isolation and protein extraction

Collagenase (C0130-100MG, Sigma-Aldrich) and hyaluronidase (H3506-100MG, Sigma-Aldrich) were dissolved in DMEM to obtain a 1 mg/ml, 50U/ml solution, respectively, which was then sterile-filtered through a 0.2 μm syringe filter (Pall Life Sciences). The hydrogel model or CAVD tissue was cut up into small pieces with a sterile razor blade and digested in the collagenase and hyaluronidase solution for 4 hours at 37 °C. The solution was mixed every 30 minutes and vortexed briefly in the end. Once finished, the digested hydrogel suspension was passed through a 40 μm cell strainer (Thermo Fisher Scientific) and the resulting solution was spun down at 1500 RPM for 5 minutes. The supernatant was aspirated, the pellet was washed in PBS and finally suspended in 20 μl RIPA Lysis and Extraction Buffer (Thermo Fisher Scientific) supplemented with PhosSTOP phosphatase inhibitor (Sigma-Aldrich) and cOmplete, protease inhibitor (Sigma-Aldrich). To maximize protein yield, acetone precipitation was used. Acetone cooled to -30°C was added to each of the samples and left to incubate for one hour at -30°C. Afterward the tubes were centrifuged for 10 minutes at 10xG after which the supernatant was aspirated. Finally, the tubes with the precipitated proteins were left to airdry until complete dryness.

### Extracellular vesicle isolation

Digestion media from hydrogel and aortic valve tissue VIC isolation, as well as culture media from 2D and 3D cultures, were used to isolate extracellular vesicles as previously described^56,84^. Briefly, digestion media and supernatant within each condition was pooled and underwent serial ultracentrifugation as follows: 2000g 10 minutes; 100kDa MWCO filtration; 10,000g 20 minutes at 4°C; 100,000g for 1 hour 4°, then washed with 1× PBS and repeated the 100,000g 1 hour spin (rotor MLA-55 in 10.4 ml 16×76 mm polycarbonate capped centrifuge tubes, Beckman Coulter 355603). The final pellet was resuspended and layered onto the top of a linear 5-step 10-30% iodixanol gradient (composed of NTE buffer and OptiPrep Density Gradient Media, Sigma-Aldrich D1556). The iodixanol gradient was then ultracentrifuged at 250,000g for 40 minutes at 4°C; fractions 1-4 were used for analysis based on findings from previous studies^56^. Pooled fractions 1-4 were topped up to a volume of 9mL with NTE buffer and underwent ultracentriguation at 100,000g for 1 hour at 4°C as above. Supernatant was discarded from each fraction, and the resultant pellets were resuspended in buffers appropriate for proteolysis or nanoparticle tracking analysis.

### Nanoparticle tracking analysis (NTA)

EV particle size and concentration was measured using NTA (Malvern Instruments, NanoSight LM10). Samples were diluted 1:500 in PBS to ∼10^9^ particles/mL. For each sample, five data collection windows (1 minute per window) were recorded during continuous injection by syringe pump with the following parameters: screen gain 1.0, 10.0 (capture, processing); camera level 9.0; detection threshold 2.0. Particle counts presented as histograms per collection per donor as well as sum normalization per donor. Data are presented as mean ± standard error, and a student’s t-test (two-tailed, unpaired).

### Mass spectrometry

For mass spectrometry samples were prepared using the Preomics iST kit (PreOmics GmbH, Planegg/Martinsried, Germany) protocol. A total of 10 μg of protein was loaded for cellular proteomes and 5 μg of protein for EV Cargo proteomes. Next, the entire protein samples were suspended in 42 μl LC-LOAD (PreOmics GmbH). An Orbitrap Exploris 480 mass spectrometer fronted with an EASY-Spray Source (heated at 45 °C), coupled to an Easy-nLC1200 HPLC pump (Thermo Scientific) was used to analyze the peptide samples. The peptides were passed through a dual column setup with an Acclaim PepMap RSLC C18 trap analytical column, 75 μm X 20 mm (pre-column), and an EASY-Spray LC column, 75 μm X 250 mm (Thermo Fisher Scientific). The analytical gradient was run at 300 nl/min from a 5% to 21% solution of Solvent B (acetonitrile/0.1 % formic acid) for 70 minutes, followed by a 21% to 30% solvent B solution for 10 minutes, and finally a 95% solvent B solution for 10 minutes. A 60K resolution was used on the Orbitrap analyzer and the top S precursor ions that were in a m/z 375-1100 scan range (60 seconds dynamic exclusion enabled) within a 3 second cycle were fragmented by high-energy collision induces dissociation. (HCD; collision energy, 26%; isolation window, 1.6 m/z; AGC target, 1.0 e4). For peptide sequencing (MS/MS), a rapid scan rate was set on the ion trap analyzer.

The SEQUEST-HT search algorithm of the Proteome Discoverer package (PD, Version 2.5) was used to run the MS/MS spectra through the Human UniProt database (101043 entries, updated January 2022) and identify the proteins present in the samples. Media digest controls were run against relevant background proteomes (bovine, porcine, c. histolyticum). Cellular and extracellular vesicle datasets were searched separately. The digestion enzyme was set to trypsin and up to two missed cleavages were allowed. Furthermore, the precursor tolerance was set to 10 ppm and the fragment tolerance window to 0.02 Da. Methionine oxidation, n-terminal acetylation was set as variable modification and cysteine carbamidomethylation as fixed modification. A peptide false discovery rate of 1.0%, which was calculated by using the PD provided Percolator, was applied to the detected peptides. Peptides that were only assigned to one given protein group and not detected in any other protein group were considered unique and used for further analyses. Additionally, a minimum of at least 2 unique peptides for each protein was required for the protein to be included in the analyses. The “Feature Mapper” was enabled in PD to identify peptide precursors that may not have been sequenced in all samples but were detected in the MS1. The chromatographic spectra were aligned while allowing for a maximum retention time shift of 10 minutes and mass tolerance of 10 ppm. Chromatographic intensities were used to establish precursor peptide abundance.

Contaminant hydrogel peptides (Fig. 3b) were excluded from analysis. The quantified proteins were exported from Proteome Discoverer (v2.5) and further analyzed using Perseus^85^. The datasets were z-score normalized and thresholded for 70% valid values across 2D, 3D, and AV conditions. Values were Log_2_ transformed for subsequent statistical analysis. Significantly differentially enriched proteins were calculated using a two-group comparison (t*-test) and a Benjamini-Hochberg false discovery rate (FDR) thresholding with a s0 value of 0.1.

### Multi-layer integration analyses

#### Comparing calcification models using LIONESS

To perform LIONESS (Linear Interpolation to Obtain Network Estimates for Single Samples), we use the proteomics data described above, with 272 proteins in EV and 2457 in cells across 22 samples. To avoid the removal of sample which have missing values in cell (∼ 5%) and EV(∼ 13%) EV proteomics,we impute the abundance using missMDA (version 1.18)^86^ implemented in R (4.2.2). missMDA considers the structure of data and performs (regularized) iterative PCA algorithm (**Supp. Fig. 7a-f))**.

First, we reduce the dimension of the cell and EV datasets and select principal components (PCs) that account for 95% of the variance in each layer (**Supp. Fig. 7c**). We then compute spearman correlation between PCs from EV and Cell datasets (**Supp. Fig. 7d**). As the network was constructed using information from all samples, it cannot independently elucidate the connections that may be linked to differences in the phenotypic properties of the input samples, such as culture and media. We then calculated sample specific networks using LIONESS implemented in R. The edge-weights of these sample-specific networks were analyzed differential correlation patterns between different calcification models (2D-OM, 2D-PM, 3D- OM,3D-PM), each compared with normal media using limma (version 3.54.2) in R (**Supp. Fig. 7e**). The top 2 most significant edges were extracted for each comparison and top 20 proteins from the loadings from these PCs were depicted in Figure 6a.

#### Multivariate analysis on Cell and EV proteome using rCCA

While LIONESS analysis identified patterns of associations between two omics layers and compared differences between calcification models, we wanted to understand the connections and dependencies between two datasets while considering the underlying structure and complexity. For that, we used Regularised Canonical Correlation Analysis (rCCA) is a multivariate approach to highlight correlations between two data sets and considers the interrelationships between variables within each dataset and explores the associations between the two datasets. We implemented rCCA from mixOmics (version 6.22.0)^87^ in R. For regularization, we used the shrinkage method and obtained exact tuned values of λ1, λ2 which better at characterized the AV samples in Canonical variate 1 and 2D from 3D and AV in Canonical variate 2.

