## Supplemental Figures for "Intracellular Proteomics and Extracellular Vesiculomics as a Metric of Disease Recapitulation in 3D Bioprinted Aortic Valve Arrays"

### Integration of Cellular and Extracelluar Vesicle Proteomics to Assess 3D Bioprinted Model of CAVD

### Supplemental Figure 1:

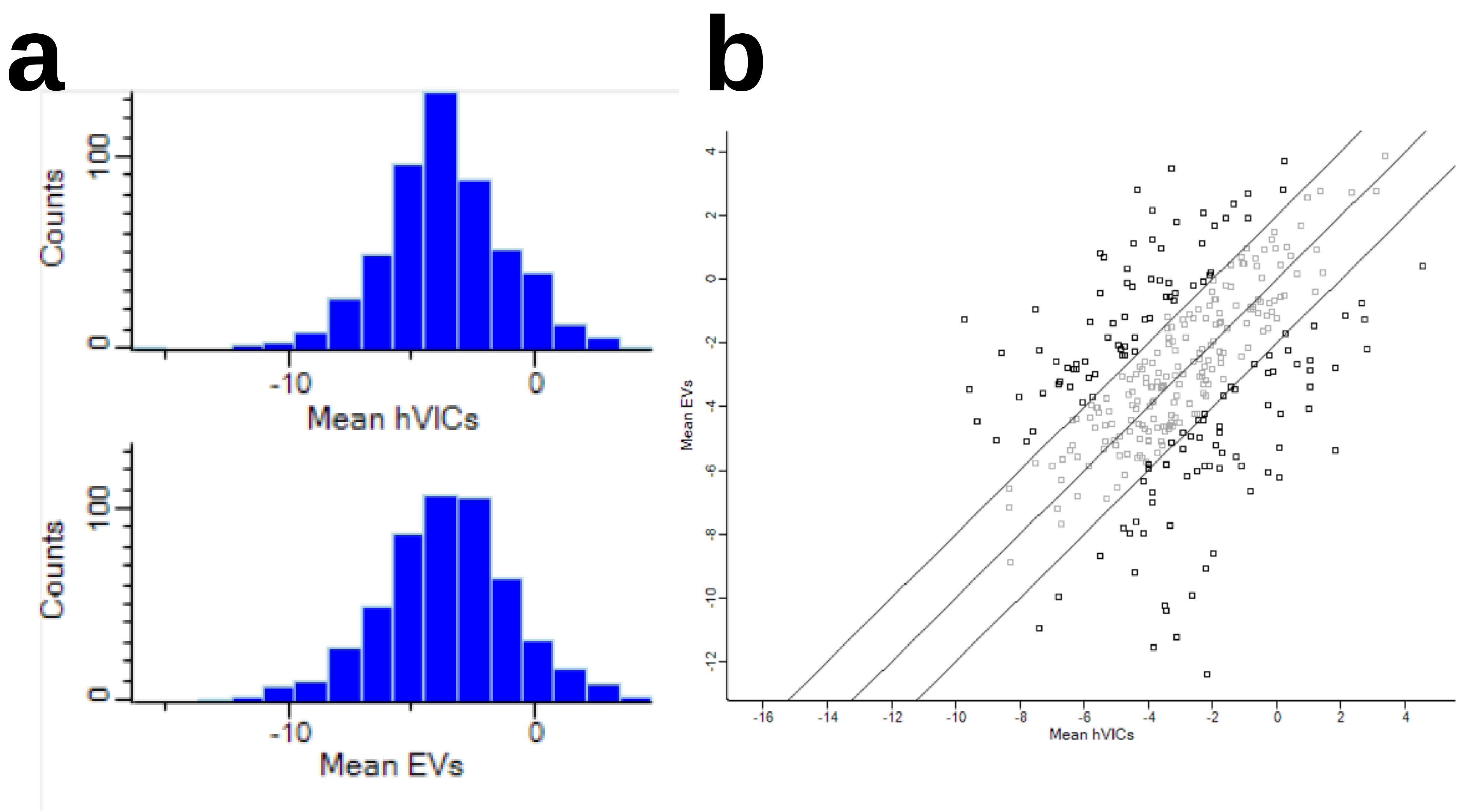


**Supplemental Figure 1: Normalization strategy for proteomics datasets**. A) Histograms show gaussians distribution of z-score normalized protein abundances for cellular and EV-derived protein datasets. B) Log2(fold change) scatterplots show, amongst proteins shared between each dataset, less than 30% are differentially enriched and are normally distributed, suggesting lack of technical variability.

### Supplemental Figure 2:

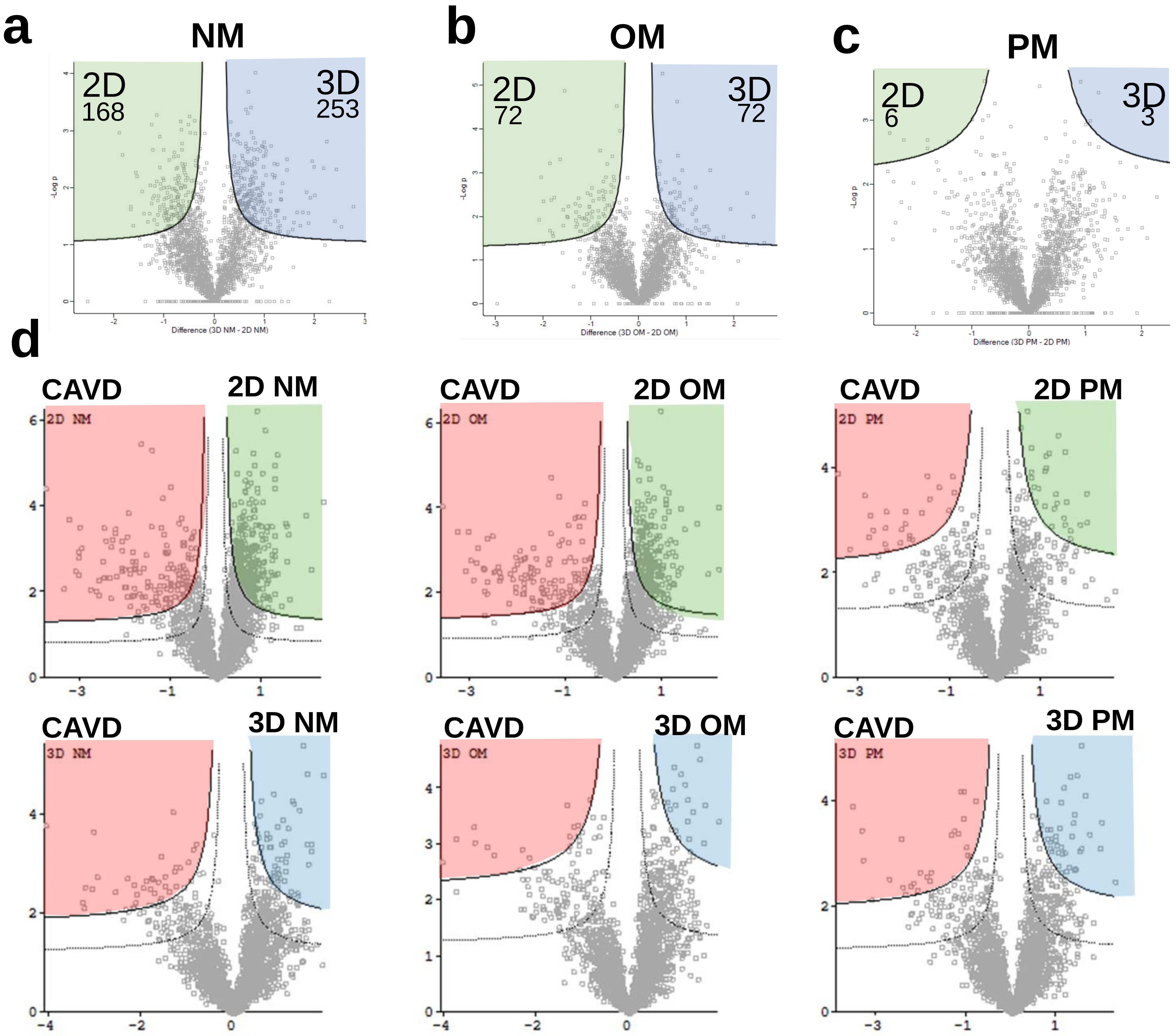


**Supplemental Figure 2:** Volcano plots of differential enrichment analysis between 2D and 3D in vitro culture models. A) NM: normal media. B) OM: osteogenic media. C) PM: pro-calcifying media. Corresponding to data in **Figure 3h-j**. D) Hawaii plot of all in vitro conditions compared to CAVD tissue. This data informs **Figure 4b-c**.

### Supplemental Figure 3:

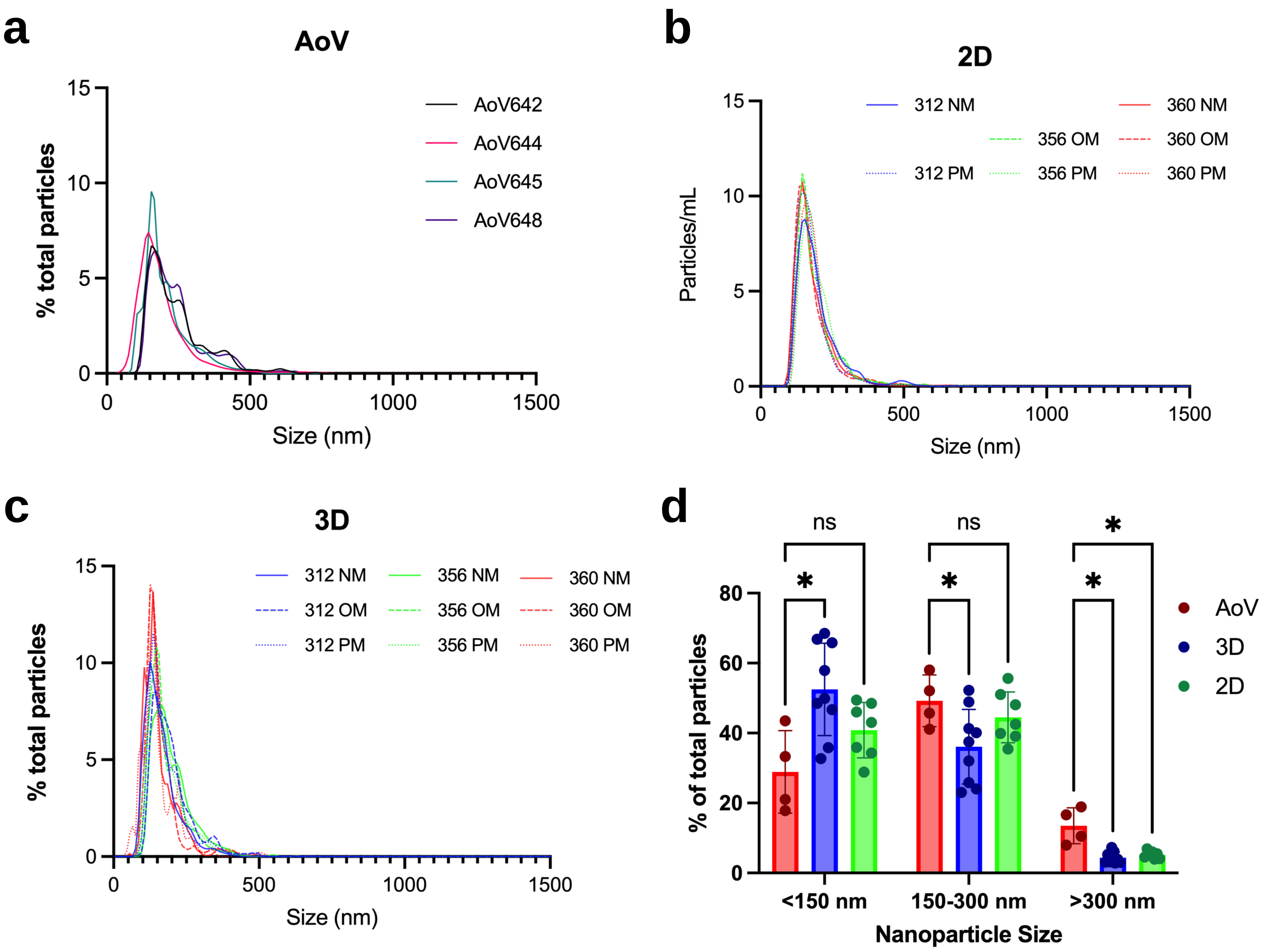


**Supplemental Figure 3: Nanoparticle tracking analysis validates extracellular vesicle size and count distribution across *in vitro* and *ex vivo* conditions. A-C)** Histograms of particle count per particle size (nm). AoV: CAVD tissue isolated EVs. 3D: 3D bioprinted hydrogel derived EVs. 2D: traditional cell culture media derived EVs. **D)** Bar chart of quantified nanoparitlce sizes for bins <150nm (small EVs), 150-300nm (large EVs) and <300nm. Fisher's LSD test against AoV *p<0.05

### Supplemental Figure 4:

a

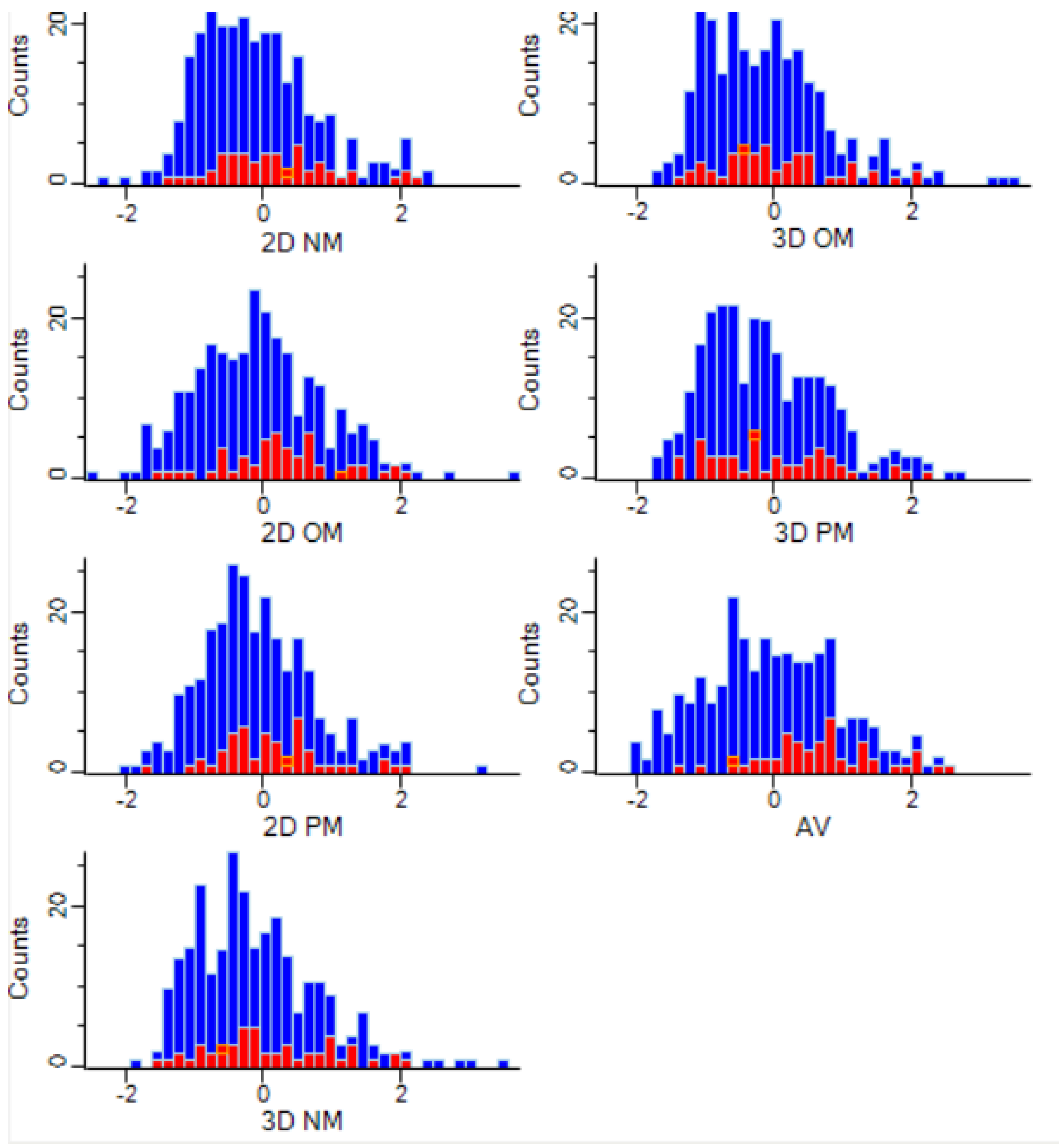


b

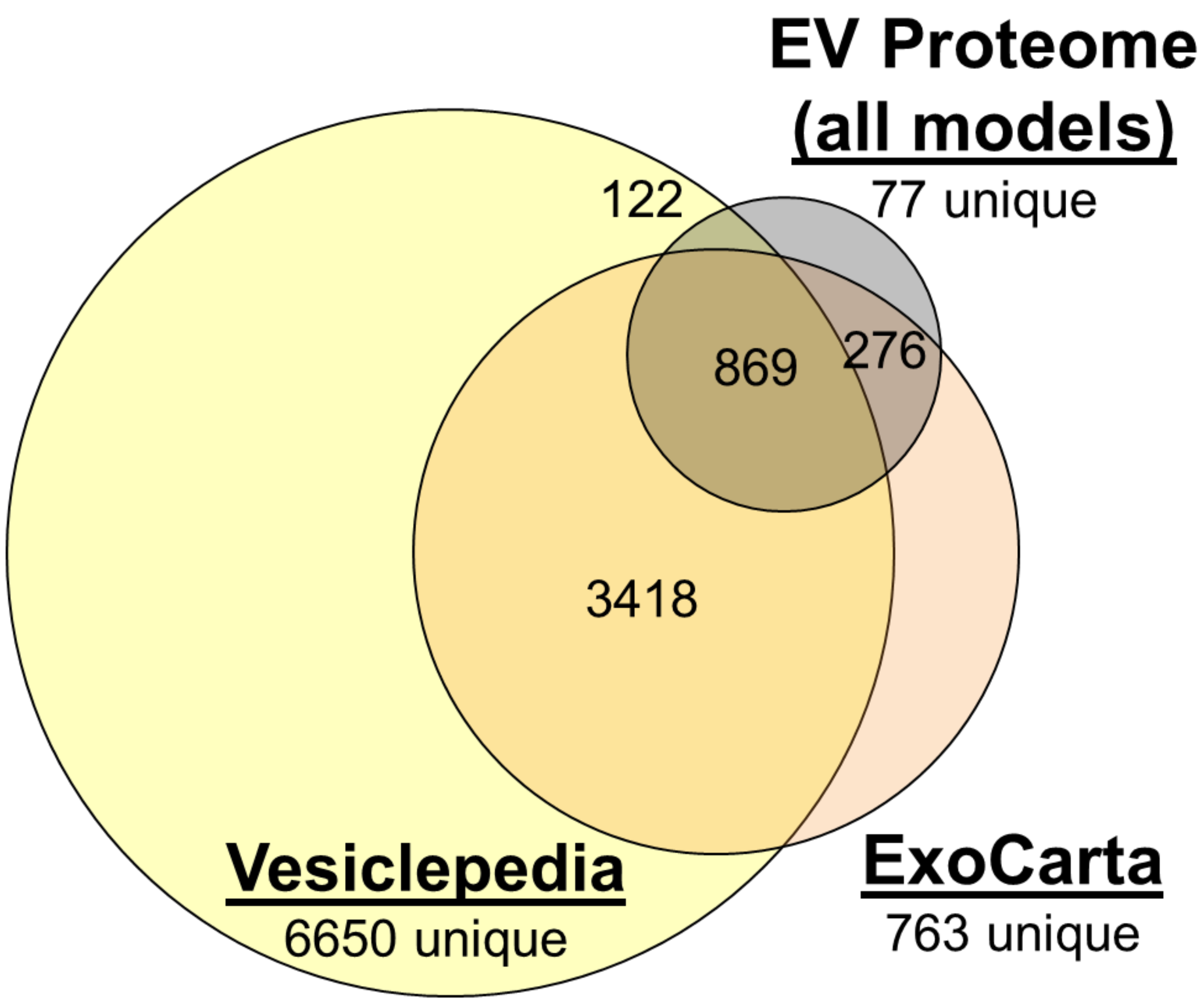


**Supplemental Figure 4: Quality control of extracellular vesicle proteomics dataset. A)** Histograms of protein abundance distribution of EV proteomics dataset for sample groups shown. Red: EV markers. Blue: all other EV-derived proteins identified. No significant change in EV marker distribution is seen relative to overall proteome. **B)** Comparison of proteins identified in this study's dataset (black) compared to public EV marker repositories such as Vesiclepedia (yellow) and ExoCarta (orange).

### Supplemental Figure 5:

**a**

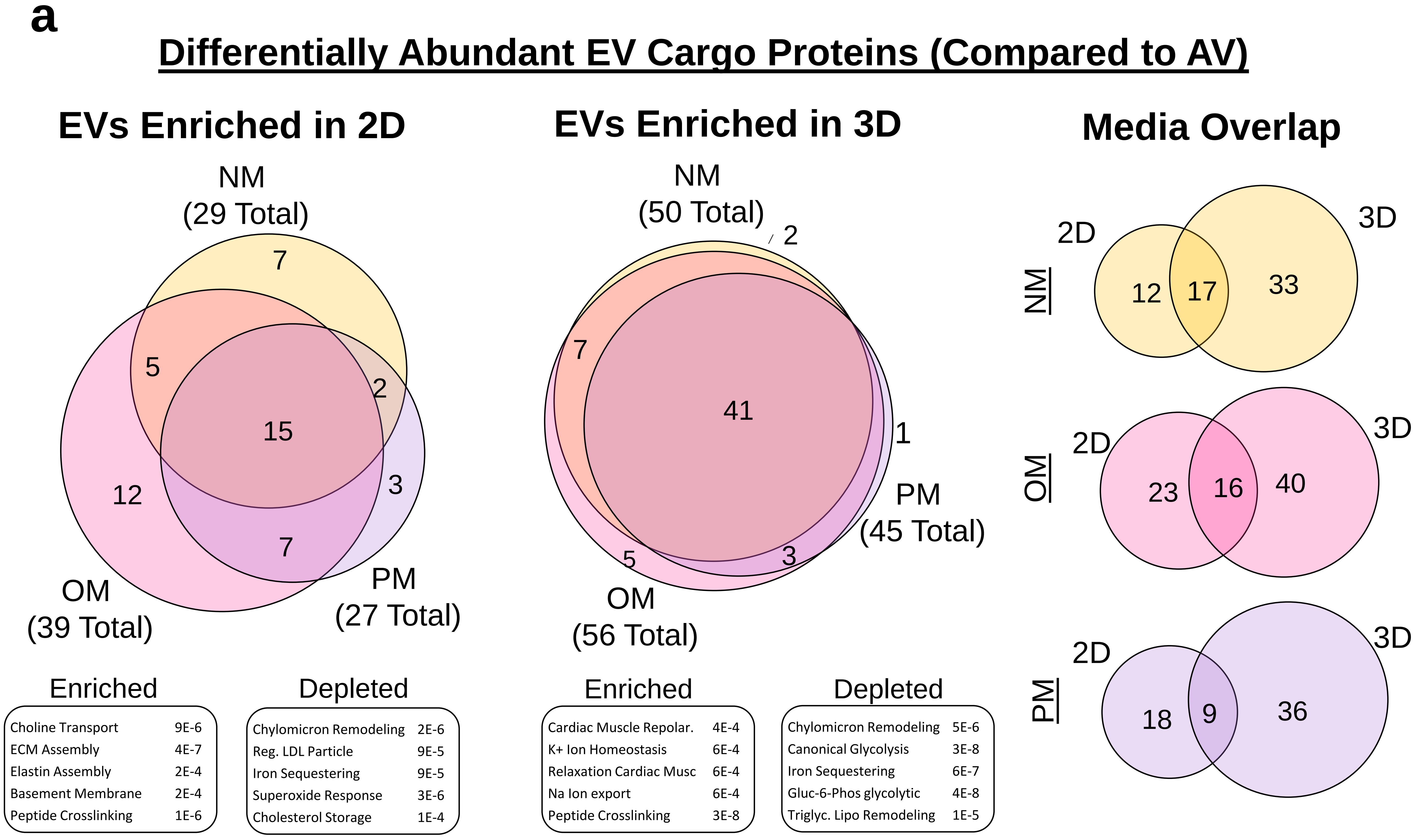


**Supplemental Figure 5: EV cargo proteins between 2D and 3D media conditions have differential disease recapitulation compared to CAVD tissue derived EVs.** Using differentially enriched cargo proteins identified in **Figure 5i**. Comparing number of proteins differentially enriched to CAVD tissue EVs; venn diagram shows number of proteins shared between NM, OM, and PM media. Gene Ontology Biological Processes are shown below 2D and 3D Venn Diagrams to described enriched and depleted terms describing the DE proteins GO Term and adjusted p-value are shown. Media overlap shows shared and unique protein numbers for each media type (NM, OM, and PM) for 2D vs 3D conditions. Few EV cargo proteins (~26%) are similarly differentially enriched in 2D and 3D conditions compared to CAVD tissue derived EV cargo.

### Supplemental Figure 6

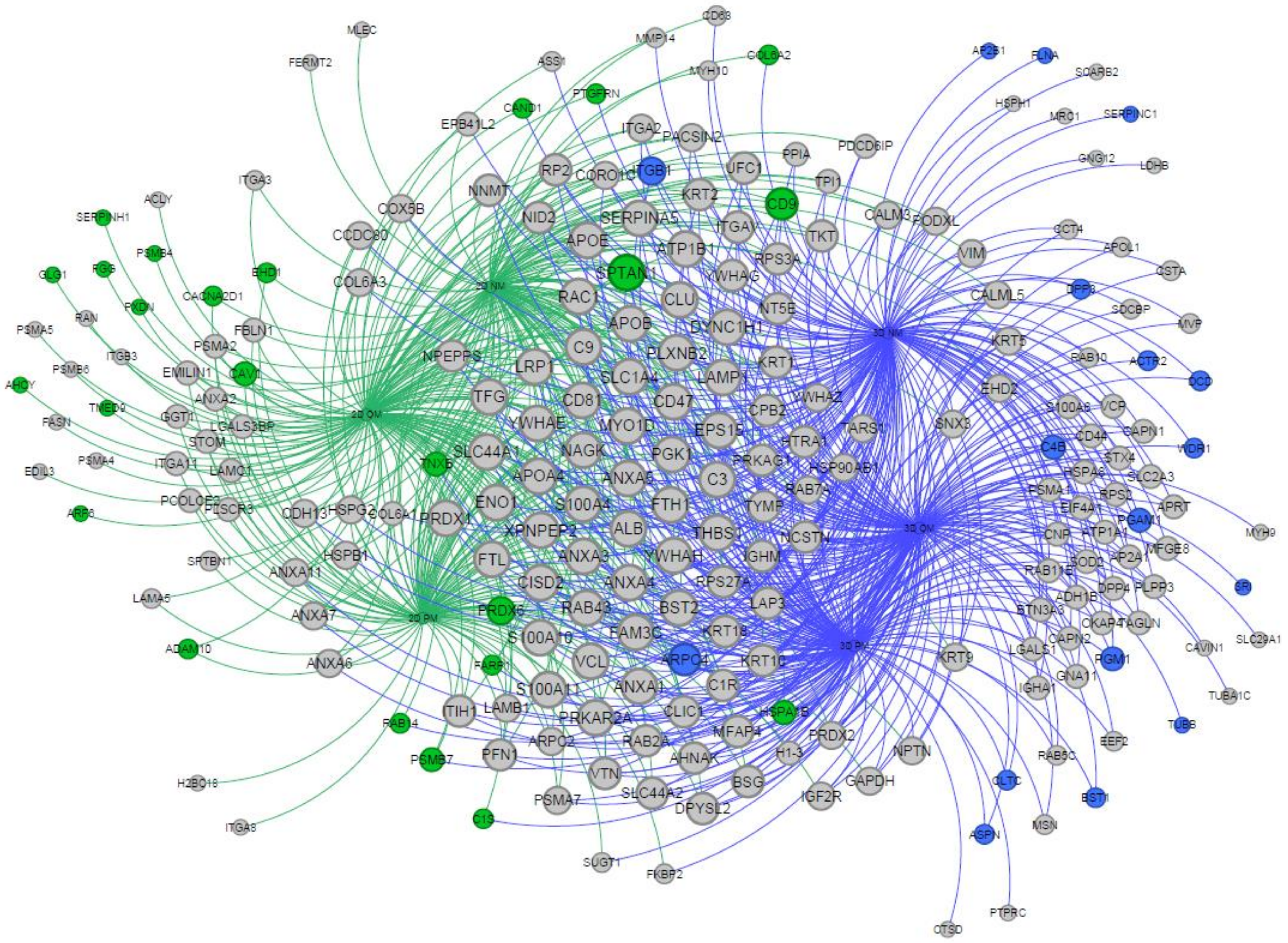


**Supplemental Figure 6. Network of top differentially enriched proteins amongst each in vitro condition compared to native CAVD tissue.** Network is exported from Perseus v2.0.0.1 Hawaii analysis (**Fig. 5i**) and formatted using Gephi v0.10.1**.** Proteins from this figure inform network in **Figure 5j**. Green = 2D unique proteins. Blue = 3D unique proteins. Grey = shared between 2D and 3D conditions as being differentially enriched compared to CAVD.

### Supplemental Figure 7:

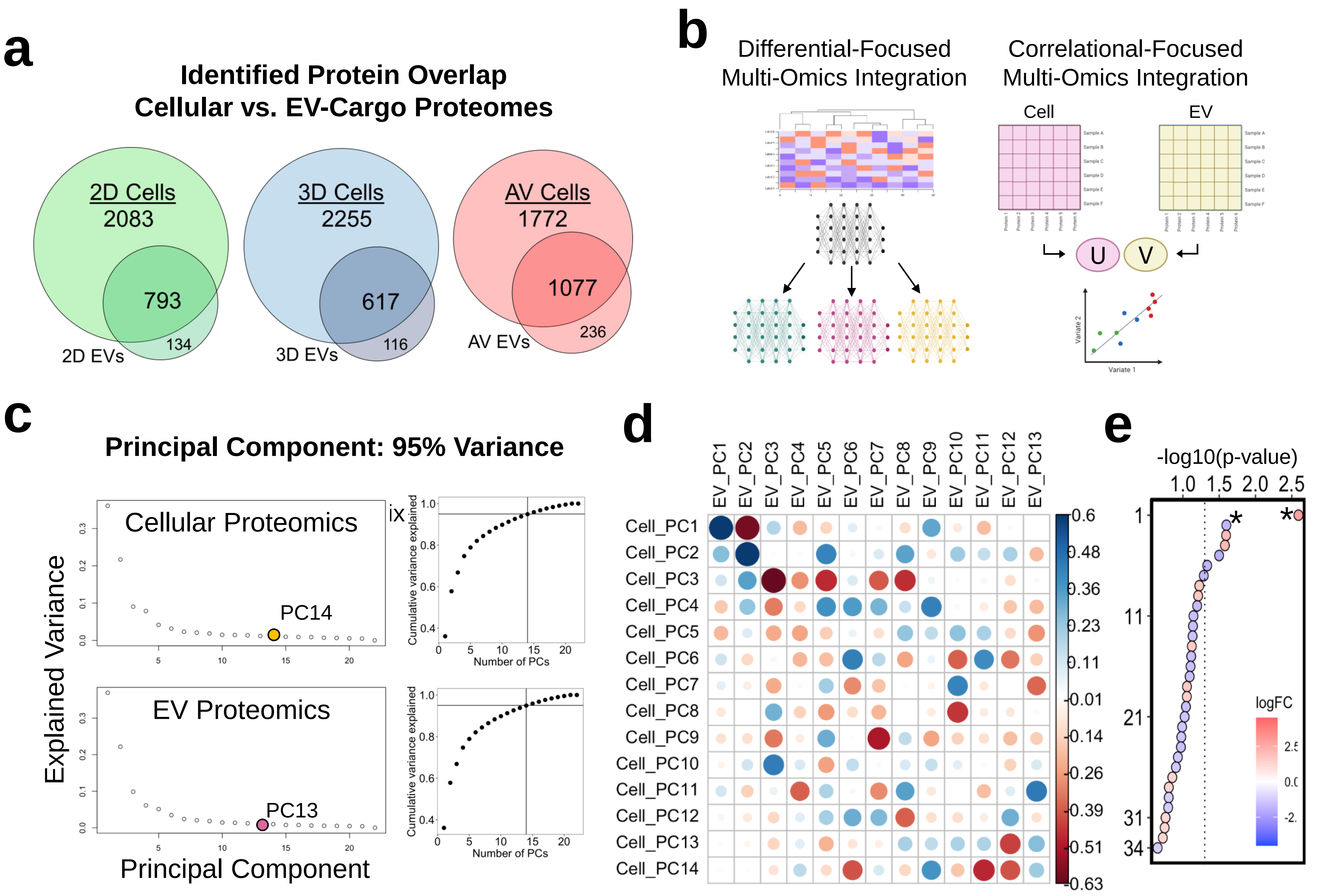


**Supplemental Figure 7: Multi-omics integration of cellular and EV cargo derived proteome reveal key drivers of CAVD pathology in 3D bioprinted model. a)** Venn diagram of number of shared and unique protein identifications between cell and EV proteomic datasets. **b)** Multi-omics integration strategy using LIONESS **(b-e)** and rCCA **(f-i)** workflows. Images modified from (Kuijjer, 2019) and (Wang, 2020). LIONESS: Linear Interpolation to Obtain Network Estimates for Single Samples. rCCA: regularized canonical correlation analysis. **b)** Distribution of the variance explained by each PC, as well as cumulative variance. The vertical and horizontal lines indicate the number of PCs which explain 95% of the variance and which were included in our network analysis. **c)** Spearman correlation across the loadings of the top 14 cellular proteome PCs and top 13 EV proteome PCs. **d)** The significance of each edge, as define from multivariate linear analysis comparing edge-weights between all groups (2D, 3D, and AV). *=top edges used for analysis.

### Supplemental Figure 8:

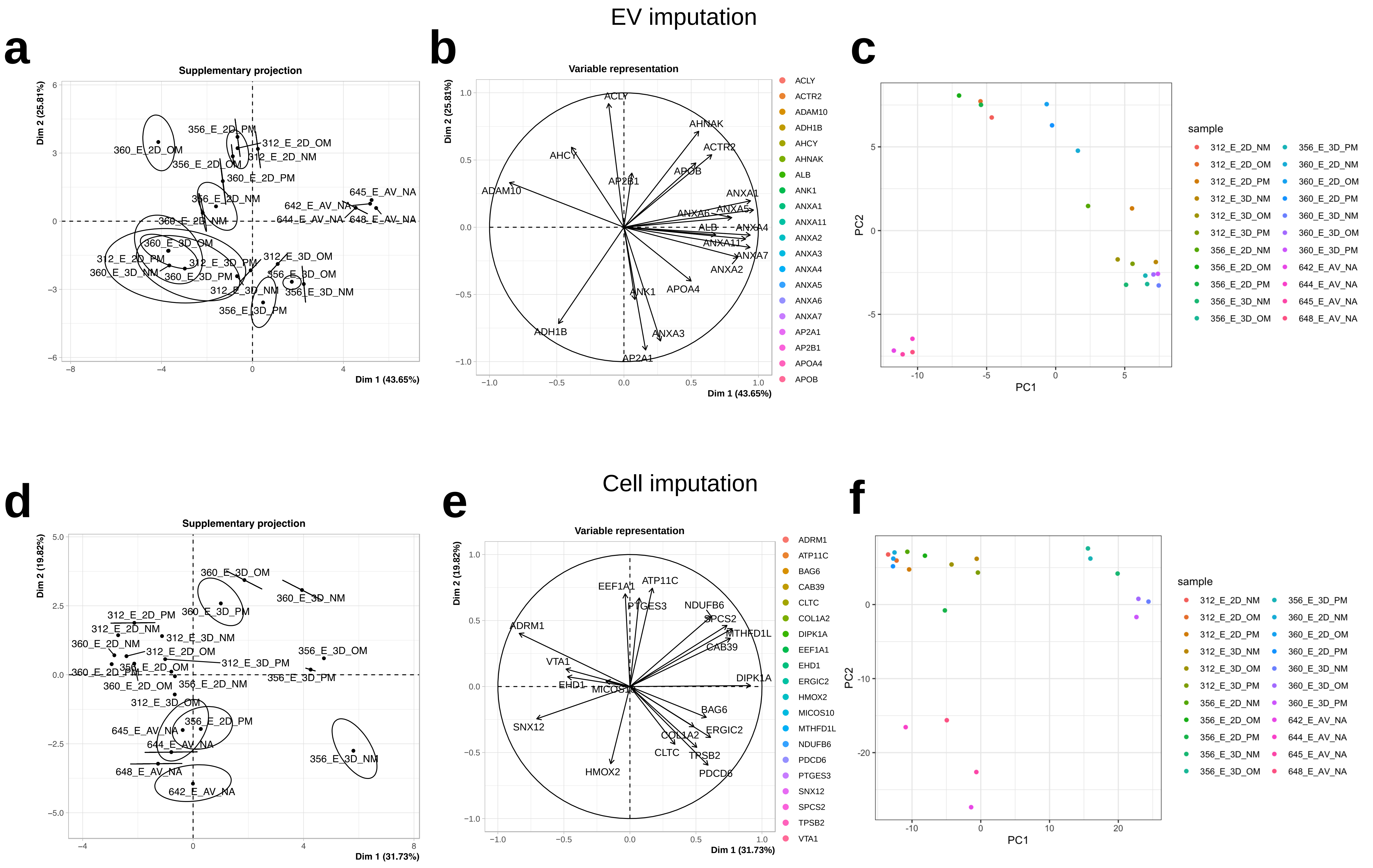


**Supplemental Figure 8: Imputation and PCA representation for LIONESS analysis.** The projection as supplementary elements of the samples and proteins of the imputed data sets for EV proteome (**A-B**) and for Cell proteome (**D-E**). **C,F**) PCA representation for EV (**C**) and Cell (**F**).
