## Supplementary material for "Intracellular Proteomics and Extracellular Vesiculomics as a Metric of Disease Recapitulation in 3D Bioprinted Aortic Valve Arrays": Figure Legends

**Figure 1: 3D bioprinting of aortic valve model**. **a)** Schematic of hybrid GelMA/HAMA hydrogel synthesis Black = gelatin, Red = hyaluronic acid, green = crosslinks, UV = photo crosslinking with 365 nm UV light at 2.5 mW/cm$^2$; **b)** Schematic illustration of the printing regime. The first printhead loaded with pluronic gel was used to print the outer ring at a pressure of 180 kPa and print speed of 6 mm/s. The second toolhead which contained the hydrogel mixture dispensed this cooled gel (20 ˚C) into the ring at a pressure of 75 kPa and 2 mm/s print speed. The hydrogel was allowed to melt and evenly spread inside the pluronic ring on top of the 37 ˚C heated printbed (not shown) after which the construct was crosslinked by the third toolhead with 365 nm UV light (UV-X) with an intensity of 2.5 mW/cm$^2$ for a total of 88 seconds (image not to scale); **c')** Acellular 3D printed GelMA/HAMA hydrogel post-pluronic ring dissolving. **c")** VIC-seeded bioprinted CAVD model. Cells seeded at a concentration of $10^7$ cells/mL. **c"')** Models were bioprinted directly into 96-well plate for immediate culture *in situ*. Scale bar = 1 mm. **d)** Heatmap of the storage modulus of the spongiosa- and fibrosa-like hydrogel. **e)** Nanoindentation experiment quantification of young's modulus (left) and storage modulus (right). **f)** Maximum intensity projections of viability stain on VICs in a fibrosa-like hydrogel at day 3 post-printing, where medium was added directly after (left) or delayed by 2 hours (right). Green = calcien AIM, Red = propidium iodide. Scale bar = 100 μm. **g)** Quantification of cell viability. Mean ± SD; ** $p < 0.01$; *** $p < 0.001$ (left); on day 1 post printing, immediate addition of media ensured consistent viability of VICs across bioprinted array.

**Figure 2: VICs cultured in fibrosa-like hydrogels in organic and inorganic phosphate media conditions maintain viability and induce calcification. a)** 3D bioprinted VICs show high viability after 14 days culture in normal (NM), organic phosphate rich osteogenic media (OM) and inorganic phosphate rich pro-calcifying media (PM) conditions. Quantification of viability assay seen in **(b)**. Negligible apoptotic staining was visualized or quantified **(b)** at the same timepoint. **c)** Representative images of Osteosense stained VIC-encapsulated fibrosa-like hydrogels cultured in NM, OM or PM for 14 days show an increased positive stain in the PM condition. White: Osteosense 680EX bisphosphonate dye **d)** the number of Osteosense positive noduli is significantly increased in the PM condition compared to NM and OM **e)** intensity of the calcific noduli was significantly higher in the PM culture condition compared to NM and OM. Mean + SD, **** $p < 0.0001$. Scale bar = 100 μm, n=3

**Figure 3: Cellular proteomics of CAVD hydrogel model reveal translational targets of modeled pathology. a)** Proteomics workflow, including models studied (2D: two dimensional VIC monoculture; 3D: 3D bioprinted VIC-seeded fibrosa-like hydrogel; AV: Fresh Tissue cell and EV isolation), media conditions (NM: normal media, OM: osteogenic media; PM: pro-calcifying media), and multi-omics analysis pipeline. **b)** Venn diagram showing overlap of identified proteins from in vitro VIC datasets compared to cell-free GelMA/HAMA hydrogel proteomics alone. GelMA/HAMA hydrogel was searched against porcine, bovine, and human databases for potential contaminant sequences. Top 3 Gene Ontology Biological Processes (GO BPs) are shown with corresponding proteins. **c)** Venn diagram showing overlap of all three CAVD model cellular proteomics datasets, highlighting unique proteins identified within each dataset. **d)** Principal Component Analysis (PCA) plot of the three cellular proteomics datasets. **e)** PCA plot of the two in vitro datasets, colored according to media treatment. **f)** Representative volcano plots informing differentially abundant proteins (#) within each dataset, without considering media treatment. **g)** Bubbleplots showing enriched GO BPs amongst each dataset. AV GOBP terms described enriched terms in *ex vivo* VIC proteome consistent with both models **h)** Bar plot of number of differentially enriched (DE) proteins amongst the *in vitro* models, by media type. **i)** LOG2(Fold Change) scatterplot of significantly differentially enriched proteins between pro-calcifying (PM) and normal media conditions, with 2D enriched proteins shown on the x-axis and 3D enriched proteins on the y-axis. Proteins similarly identified between each dataset fall along the outer quandrants. . **j)** LOG2(Fold Change) scatterplot of significantly differentially enriched proteins between osteogenic (OM) and normal media conditions. FDR 0.05; $S_0$ 0.1.

**Figure 4: 3D Bioprinted model cellular proteome best recapitulates CAVD cellular pathology in calcifying conditions. a)** Multi-scatterplot showing LOG2(Fold Change) abundances of proteins between all media and model conditions. Top-right half shows density plots, while bottom-left half shows correlation coefficients of corresponding scatterplots. **b)** Bar chart of number of differentially enriched proteins per media condition, comparing AV and 2D conditions. **c)** Bar chart of number of differentially enriched proteins per media condition, comparing AV and 3D conditions. **(b-c)** Quantified from post-hoc corrected Hawaii analysis (**Supplemental Figure 2**). **d,g)** Heatmap of thresholded differentially abundant proteins across all model and media conditions. Heatmap cluster trend plots highlight isolated proteins that are similarly expressed between 2D OM vs AV and 2D PM vs AV conditions. Top GO BP are highlighted with corresponding adjusted p-values. **e-i)** GOBP-Protein network highlighting key proteins driving similarities between 2D PM and CAVD (decreased, **e**; increased **f**) and 3D PM and AV (decreased, **h**; increased **i**). As discussed in text, spontaneous calcification occurred in 3D model only, NM conditions in this model showed trending protein abundances as well. ANOVA $q \leq 0.05$ Tukey HSD; FDR 0.05; $S_0$ 0.1.

**Figure 5: Extracellular vesicle proteomics identifies matrix-dependent cargo loading and ubiquitous AV associated cargo a)** Extracellular vesicle proteomics identified over 1300 proteins across all three models and media conditions, with 26 EV markers identified. **b)** Unbiased PCA clustering of all three EV datasets, colored by model. **c-d)** Unbiased PCA cluster of 2D **(c)** and 3D **(d)** EV proteomic datasets, colored by media condition. **e)** Quantification of number of differentially expressed EV cargo proteins between in vitro models for NM, OM, and PM media conditions. **f-h)** Volcano plots of data quantified in (d) for NM **(f)**, OM **(g)**, and PM **(h)**, Only significant DE proteins with LOF2FC >1 have legend callouts (**f-h**). **i)** Hawaii plot of EV proteomic in vitro datasets, compared against CAVD EV cargo proteome, corrected for multiple comparisons. Key dysregulated sEV and lEV markers are highlighted. sEV: small EVs, orange; lEV: large EVs, green. Solid Line: FDR 0.01%, $S_0$ 0.1; Dotted Line: FDR 0.05%, $S_0$ 0.1. **j)** Network showing top 5 gene ontology biological processes and their corresponding proteins, curated from all differentially expressed proteins in panel **(i)**. 1. Extracellular Matrix Organization

(GO:0030198), 2. Extracellular Structure Organization (GO:0043062). 3. Neutrophil Mediated Immunity (GO:0002446), 4. Neutrophil degranulation (GO:0043312). 5. Neutrophil Activation Involved in Immune Response (GO:0002283). **k)** Top GO BP of *in vitro* derived EV cargo proteins correlated with CAVD tissue derived EV Cargo proteins, with corresponding adjusted p-value (**Supplemental Table 3**).

**Figure 6: Multi-omics integration of cellular and EV cargo derived proteome reveal key drivers of CAVD pathology in 3D bioprinted model. a)** Integrated network of top proteins from EV and cellular proteomes (ranked) that drive differential abundance between shown cases (3D PM, 3D OM, 2D PM, and 2D OM) vs NM conditions. 2 Cell and 2 EV Principal Components (PCs) were chosen with their respective top 20 proteins, ranked. Additional analysis informing this network can be seen in **Supplemental Figure 6**. **b)** Canonical correlation scatterplot of integrated cell and EV proteomics data. NM media conditions were excluded from rCCA analysis (**b-e**) to focus on correlations driving calcification. Canonical Variate 1 best describes the correlation between 3D and 2D, while Canonical Variate 2 best describes the correlation between 3D and AV. **c)** Correlation circle plot of proteins, corresponding to variants in **(b)**. Threshold 0.8. **d)** Clustered Image Map (CIM) showing correlation between EV and Cell derived proteins used for integration. Red (or blue) indicates high positive (or negative) correlation, respectively. **e)** Relevance network obtained rCCA analysis significant proteins. Red (or green) indicate high positive (or negative) correlation, respectively. Node size represents betweenness centrality.
